# A sense-antisense RNA interaction promotes breast cancer metastasis via regulation of NQO1 expression

**DOI:** 10.1101/2021.10.08.463652

**Authors:** Bruce Culbertson, Kristle Garcia, Daniel Markett, Hosseinali Asgharian, Li Chen, Lisa Fish, Albertas Navickas, Johnny Yu, Brian Woo, Scott Nanda, Joshua Rabinowitz, Hani Goodarzi

## Abstract

Antisense RNAs are ubiquitous in human cells, yet the role that they play in healthy and diseased states remains largely unexplored. Here, we developed a computational framework to catalog and profile antisense RNAs and applied it to poorly and highly metastatic breast cancer cell lines. We identified one antisense RNA that plays a functional role in driving breast cancer progression by upregulating the redox enzyme NQO1, and hence named NQO1-antisense RNA or NQO1-AS. This upregulation occurs via a stabilizing interaction between NQO1-AS and its complementary region in the 3’UTR of NQO1 mRNA. By increasing expression of NQO1 protein, breast cancer cells are able to tolerate higher levels of oxidative stress, enabling them to colonize the lung. During this process the cancer cells become dependent on NQO1 to protect them from ferroptosis. We have shown that this dependence can be exploited therapeutically in xenograft models of metastasis. Together, our findings establish a previously unknown role for NQO1-AS in the progression of breast cancer by serving as a post-transcriptional regulator of RNA processing and decay for its sense mRNA.

## Introduction

Metastasis is a disease of disordered gene expression^1,2^. During cancer progression, gene expression patterns change to promote uninhibited cell growth and spread. Understanding the cellular pathways that underlie gene expression changes is a necessary step towards developing therapies that target metastatic progression. It has become increasingly clear that cancer cells often co-opt post-transcriptional regulatory mechanisms to achieve pathologic expression of cellular pathways that impact metastasis^3–5^. While RNA-binding proteins^6,7^ and microRNAs^8^ have been the focus of many studies on cancer progression, it has been hypothesized that interactions between RNA transcripts may also play a regulatory role^9–11^. However, the extent to which RNA-RNA interactions influence gene expression and regulate cell physiology and human disease remains unknown.

Antisense RNAs, a major class of non-coding RNAs, have great regulatory potential, as they readily form duplexes with RNAs transcribed from their complementary sense strands. Additionally, they are ubiquitous within human cells. It has been estimated that ∼30% of human protein coding genes have a corresponding antisense RNA^12^. Yet, while antisense RNAs were discovered decades ago, little is known about their regulatory functions in the cell. Specific members of this family have been shown to impact gene expression through a number of molecular mechanisms, including DNA methylation^13^, chromatin modification^14^, and RNA degradation^15^. A number of antisense RNAs have also been associated with tumorigenesis, and a select few have been shown to play a functional role in cancer progression^16^. Still, the extent to which antisense RNAs contribute to the regulation of gene expression in cancer, and the mechanisms by which this happens, remains poorly understood.

Given their ubiquity and their potential regulatory role, an unbiased and systematic study of antisense RNA in cancer is needed. To address this problem, we developed an integrative experimental and computational pipeline to catalog and profile antisense RNAs and applied it to an established xenograft model of breast cancer metastasis. Based on systematic analyses of this dataset, we identified an antisense RNA whose upregulation promotes breast cancer metastasis. This RNA is complementary to a portion of the 3’UTR of *NQO1* (NADPH quinone dehydrogenase 1), and we have therefore named it NQO1-AS. By binding directly to its complementary region, NQO1-AS stabilizes the NQO1 mRNA, upregulating NQO1 protein, a protein that protects cells against oxidative stress. Upregulating NQO1-AS therefore enables breast cancer cells to alter their metabolic profile to promote resistance to oxidative damage and to ferroptosis. During metastasis to the lung, cancer cells become dependent on this NQO1-pathway, and we show that downregulation of either NQO1 or NQO1-AS significantly decreases metastatic burden in *in vivo* mouse models. Single cell RNA sequencing experiments revealed a high degree of heterogeneity in NQO1 expression in cancer cell populations. This observation led us to test the addition of a ferroptosis inducing agent to an existing drug regimen to simultaneously target cells with both high and low NQO1 expression. This combination therapy was effective in a mouse model, and suggests a novel strategy for treating metastatic breast cancer.

## Results

### Discovery and annotation of antisense RNAs occluded by sense transcripts

We recently developed IRIS (identification of RNA antisense species), a computational pipeline that integrates data from RNA-seq, global run-on sequencing (GRO-seq), and RNA polymerase II ChIP-seq to identify and quantify antisense RNAs. We first performed a global run-on assay (GRO-seq) in MDA-MB-231 breast cancer cells to capture the footprint of transcriptionally active RNA polymerase II (RNAPII) across the transcriptome. We used the resulting data, which is stranded, to ask whether there are actively transcribed loci that show higher coverage of the antisense strand than expected by chance. For this, we used a sliding window of 500nt with a 250nt step across the transcriptome to compare sense and antisense read coverage. We used a statistical framework built on logistic regression to calculate a p-value for antisense transcription activity as measured by the number log-ratio of antisense to sense reads in each window relative to the background (a quantity we have named logASR or log of antisense to sense ratio; schematized in Fig S1a). We used the sequences with significantly positive logASR values (logASR >0.5; FDR-adjusted *p*-value <0.01) to annotate parts of the transcriptome that show significant antisense RNA signal (Fig S1b). We also generated a set of negative annotation as a control group (logASR <0; FDR-adjusted *p*-value >0.5) for the subsequent steps. In addition to the significant antisense strand bias captured by GRO-seq, we expect true antisense RNA transcription to coincide with RNAPII binding. To determine the level of RNAPII binding in our regions of interest we took advantage of POLR2A ChIP-seq datasets from ENCODE. We used our negative control set to identify a threshold above which there is strong evidence of RNAPII activity across many cell lines (Fig S1c). We chose the median plus one interquartile range to define this threshold, and, as shown in Fig S1d, we identified ∼300 loci with strong evidence for antisense RNA transcription. Finally, as an additional filter, we performed stranded RNA-seq on MDA-MB-231 cells and their highly lung metastatic MDA-LM2^1^ and used the resulting dataset to calculate logASR values for these 300 regions (Fig S1d). We selected logASR of 1.0 and adjusted *p*-value of 1e-5 for our final annotation of 262 antisense RNAs expressed in the MDA-231 background. Of these, ∼20 overlap previously annotated antisense RNAs and/or pseudogenes and the remainder were not previously reported.

We then asked whether any of these antisense RNAs are associated with the increased metastatic capacity of MDA-LM2 cells. Using our RNA-seq data from the parental MDA-MB-231 cells and their highly metastatic derivative (MDA-LM2), we performed differential gene expression analysis focused on our annotated antisense RNA species. As shown in Fig 1a, our analysis revealed three antisense RNAs that were significantly upregulated in the highly metastatic line. Among these, we chose to focus on one uncharacterized RNA that is transcribed from the antisense strand of the gene NQO1. This antisense RNA, which is annotated as CTD-2033A16.1, overlaps the 3’UTR of NQO1, here we named it NQO1-AS. While NQO1 and NQO1-AS are transcribed from distinct promoters close to 20kb apart, we observed a similar up-regulation in NQO1 expression in MDA-LM2 cells (Fig 1b-d, Fig S1e). This simultaneous up-regulation of NQO1 and NQO-AS in the highly metastatic MDA-LM2 cell line raised the possibility of their functional role in breast cancer progression.

**Figure 1.**
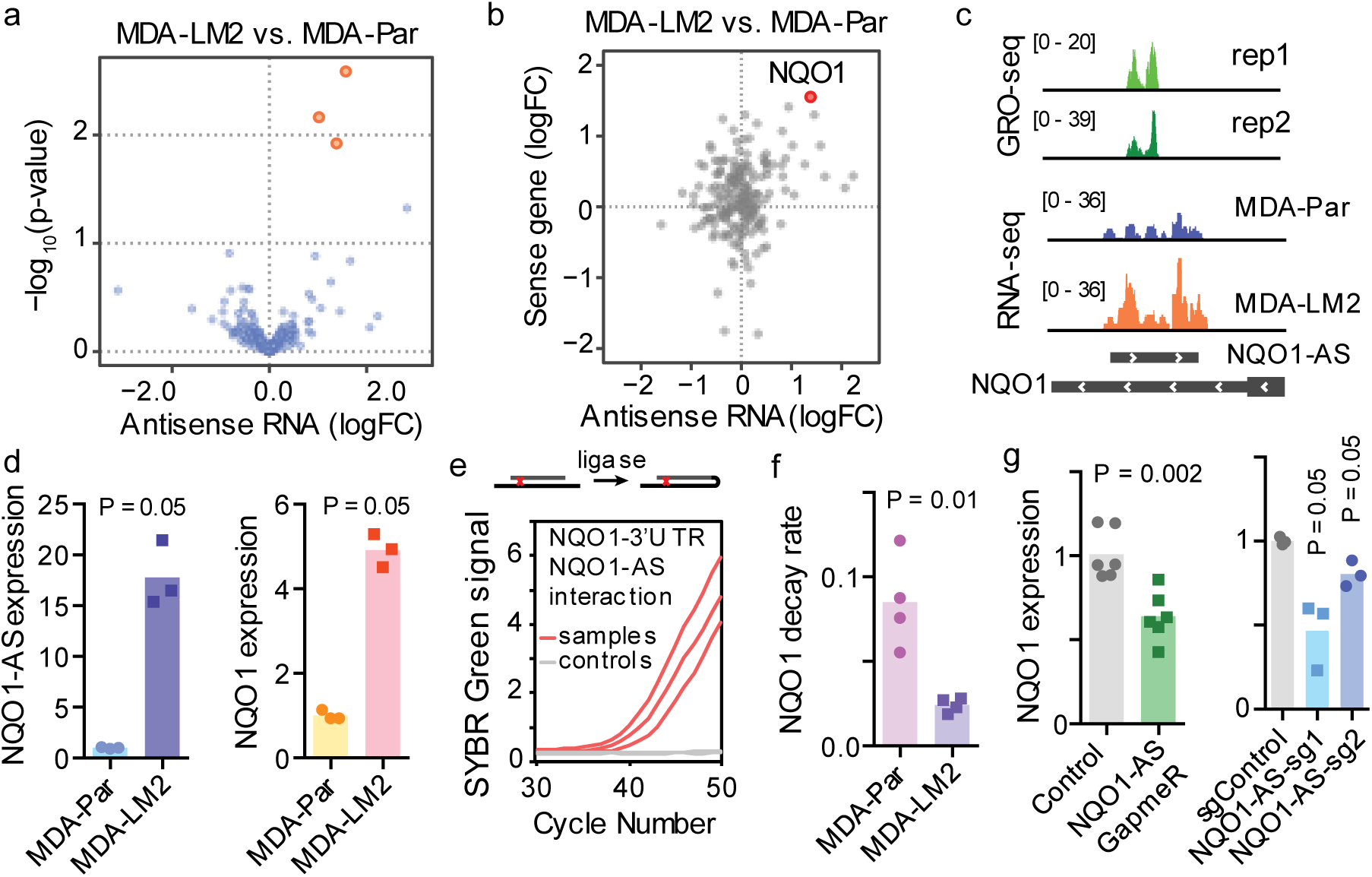
NQO1-AS interacts with the NQO1 3’UTR and regulates NQO1 expression. (a) Volcano plot showing distribution of antisense RNAs detected by RNA-seq of MDA-MB-231 (MDA-Par) and MDA-LM2 cell lines. (b) Plot showing correlation between the fold-change in MDA-LM2 compared to MDA-Par lines of antisense RNAs with their corresponding sense RNAs. NQO1 indicated in red. (c) Tracks showing global run-on sequencing in MDA-LM2 cells, along with RNA-seq in MDA-LM2 and MDA-Par lines. (d) qRT-PCR of NQO1-AS and NQO1 RNA levels in MDA-Par and MDA-LM2 lines. N = 3. (e) Psoralen-mediated RNA crosslinking followed by ligation was performed. qRT-PCR was then used to detect interactions between the NQO1 sense 3’UTR with NQO1-AS RNA. Samples without the proximity ligation step were used as controls. (f) The relative NQO1 decay rate in MDA-Par and MDA-LM2 lines from our metabolic pulse-chase labeling of these lines^3^. N = 4. (g) Relative NQO1 mRNA levels were measured using qRT-PCR in MDA-LM2 cells with gapmer-mediated NQO1-AS knockdown and with CRISPRi-mediated NQO1-AS downregulation. N = 3. All *P* values were calculated using one-tailed Mann-Whitney *U* tests.

### NQO1-AS binds and stabilizes the NQO1 sense transcript

To assess the functional relationship between NQO1-AS and NQO1, we asked if we could detect a direct interaction between the two RNA species. For this, we used psoralen crosslinking followed by nuclease digestion and proximity RNA ligation^17^. In this assay, the presence of ligase-induced artificial junctions, which can be detected using RT-PCR, revealed in vivo duplex formation between the NQO1 sense and antisense RNAs (Fig 1e). We next asked whether the upregulation of NQO1 in MDA-LM2 cells was due to increased transcription or mRNA stabilization. Whole-genome RNA stability measurements in MDA-Par and MDA-LM2 cells^3^ revealed that NQO1 mRNA is significantly stabilized (∼4-fold) in MDA-LM2 cells relative to their poorly metastatic parental line (Fig 1f). NQO1 transcription rate, as measured by pre-mRNA levels, was moderately elevated in MDA-LM2 cells (Fig S1f), but this elevation is insufficient to account for the degree of NQO1 upregulation. These results indicate that the increased levels of NQO1 mRNA in the highly metastatic cell lines is in part due to post-transcriptional stabilization and cannot be explained by transcriptional activation alone. Given that NQO1-AS is complementary to the 3’UTR of NQO1 mRNA, and psoralen crosslinking data showed an interaction between the two RNA species, we hypothesized that NQO1-AS binds and stabilizes its sense counterpart. To test this hypothesis, we used locked nucleic acid (LNA) gapmers for targeted knockdown of NQO1-AS in MDA-LM2 cells (Fig S1g). We observed that upon gapmer-induced two-fold knockdown of NQO1-AS, NQO1 mRNA levels decreased by a similar magnitude (Fig 1g). To verify this result, we repeated this experiment using CRISPRi to knock down NQO1-AS (Fig S1g). Although NQO1 pre-mRNA levels were unaffected (Fig S1h), we again saw a decrease in NQO1 mRNA levels comparable to the degree of NQO1-AS knockdown (Fig 1g).

### NQO1-AS binding modulates polyA site selection by masking HNRNPC binding sites

Having established that NQO1-AS stabilizes NQO1 mRNA by binding to its 3’UTR, we next sought to understand the mechanism by which this stabilization occurs. We hypothesized that NQO1-AS binding masks *cis*-regulatory elements targeted by other destabilizing factors. To identify these elements, we performed a sequence analysis of the 3’UTR region complementary to NQO1-AS by systematically searching for motifs over-represented in the NQO1 3’UTR relative to scrambled control sequences. We did not identify any strong recognition sites for known miRNAs among the discovered motifs; however, we observed a significant enrichment of uridine-rich motifs in this region (19 occurrences in the selected NQO1 3’UTR region compared to fewer than a median of five for the scrambled controls) (Fig S2a). To identify the specific RNA-binding protein (RBP) that binds these elements, we used publicly available cross-linking immunoprecipitation followed by RNA sequencing (CLIP-seq) datasets: CLIPdb^18^ and ENCODE eCLIP^19^, and our own CLIP-seq datasets^3,4,7,20^. We observed that the U-rich motifs in the NQO1 3’UTR were most similar to C IP-seq-derived HNRNPC binding sites (Fig S2b). These sites were defined using HNRNPC iCLIP data, which finds RBP binding sites across the transcriptome at nucleotide resolution^21^. Consistent with this, we identified five significant HNRNPC binding sites on the NQO1 3’UTR, and we observed that these binding sites correspond to U-rich motifs (Fig 2a). This observation provided evidence that HNRNPC directly interacts with the NQO1 3’UTR in vivo. Consistent with HNRNPC acting as a post-transcriptional regulator of NQO1 expression, we observed a negative correlation between HNRNPC and NQO1 levels in multiple independent datasets^22^ (Fig 2b). Furthermore, we analyzed RNA-seq data from HNRNPC knockdown cells^23^, and observed a significant increase in the expression of NQO1 upon knockdown of HNRNPC (Fig S2c). To test our hypothesis that NQO1-AS binding masks HNRNPC binding sites, we performed HNRNPC CLIP-qPCR in NQO1-AS knockdown and control cells. As expected, we observed significantly higher binding of HNRNPC to the NQO1 3’UTR upon NQO1-AS knockdown (Fig 2c).

**Figure 2.**
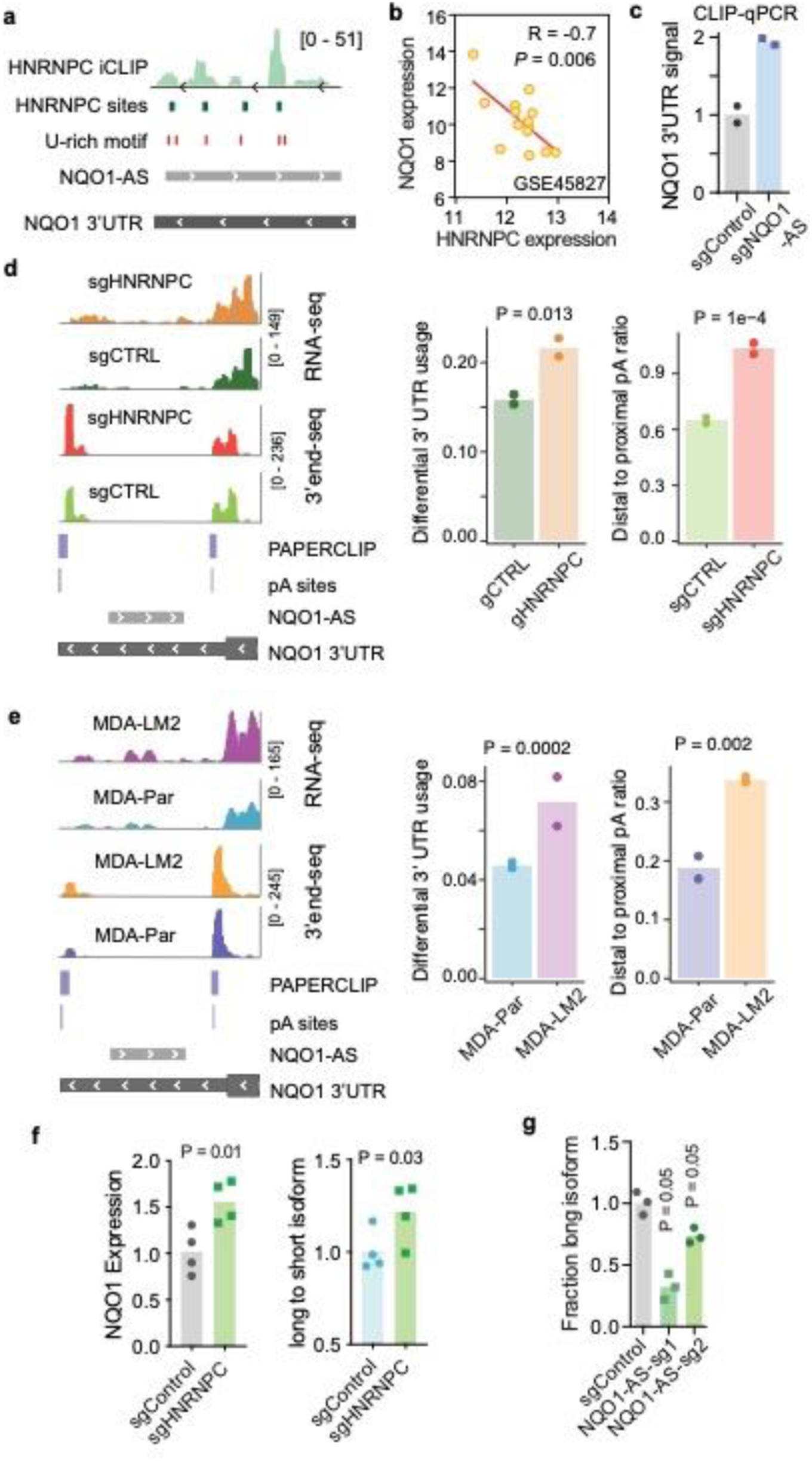
NQO1-AS masking of HNRNPC binding sites modulates polyA site selection. (a) Tracks showing the relationship between HNRNPC iCLIP data, known HNRNPC binding sites, and U-rich motifs along the NQO1 3’UTR. (b) Plot showing the correlation between NQO1 and HNRNPC expression in multiple independent datasets. (c) CLIP-qPCR with immunoprecipitation of HNRNPC in NQO1-AS knockdown and control LM2 cells. qPCR primers are targeting the NQO1 3’UTR. N = 2. (d) Read distribution of RNA-seq and 3’end-seq in MDA-Par cells with and without knockdown of HNRNPC, along with bar graph representation of differential 3’UTR usage and distal to proximal poly(A) ratio. N = 2. (e) Read distribution of RNA-seq and 3’end-seq in MDA-Par and MDA-LM2 cells, along with bar graph representation of differential 3’UTR usage and distal to proximal poly(A) ratio. N = 2. (f) qPCR showing NQO1 expression and long to short isoform ratios in HNRNPC knockdown and control MDA-LM2 cells. N = 4. (g) qPCR showing the fraction of long NQO1 isoform present in NQO1-AS knockdown and control MDA-LM2 cells. N = 3.

The above results are consistent with a model where NQO1 mRNA is stabilized when its direct interaction with HNRNPC is abolished. The underlying molecular mechanism through which HNRNPC may decrease NQO1 stability is unknown. Recently, HNRNPC has been shown to play a role in regulating transcriptome-wide poly(A) site selection^24–26^. We therefore hypothesized that HNRNPC might alter the stability of NQO1 mRNA by a similar mechanism. Our analyses revealed that the NQO1 3’UTR contains two canonical polyadenylation sites (Fig S2d). These sites were also present in an experimentally derived poly(A) site dataset, poly(A) binding protein-mediated mRNA 3’-end retrieval by crosslinking immunoprecipitation (PAPERCLIP)^27^. To address this question, we analyzed the TCGA-BRCA dataset to determine if there was a difference in the ratio of the long and short NQO1 isoforms (resulting from proximal or distal poly(A) site selection) in cells with high or low HNRNPC expression. Consistent with our hypothesis, we found that there was an increased long to short NQO1 isoform ratio in cells with low HNRNPC compared to high HNRNPC levels (Fig S2e). Next, we asked if there was a correlation between long to short NQO1 isoform ratio and NQO1-AS expression, and we found that higher NQO1-AS expression was correlated with a greater long to short NQO1 isoform ratio (Fig S2f). To test these results experimentally, we performed RNA-seq and 3’end-seq in MDA-Par cells with and without knockdown of HNRNPC^26^ (Fig 2d). Consistently, we found that HNRNPC knockdown cells had more RNA-seq reads in the portion of the NQO1 3’UTR distal to the proximal poly A site than did the controls. Additionally, 3’end-seq showed significantly higher usage of the distal poly(A) site in the HNRNPC knockdown cells (Fig 2d). These results suggest that decreasing HNRNPC expression causes an increase in the proportion of the long isoform of NQO1 mRNA, consistent with our analysis in the TCGA-BRCA dataset. Next, to determine if the higher expression of NQO1-AS in MDA-LM2 cells compared to MDA-Par cells results in greater usage of the distal poly(A) site we performed RNA-seq and 3’end-seq in these cell lines^26^ (Fig 2e). We found that MDA-LM2 cells had an increase in RNA-seq reads 3’ of the proximal poly(A) site, and there was increased use of the distal site in the 3’end-seq data. These results support a model where HNRNPC binding promotes use of the proximal poly(A) site, resulting in an increase in the NQO1 isoform with a truncated 3’UTR. When NQO1-AS interrupts this interaction, the distal poly(A) site is favored and the longer isoform is favored. To further test this model, we performed RT-qPCR using isoform-specific primers in HNRNPC knockdown and control MDA-Par cells. Upon knockdown of HNRNPC, we observed an increase in NQO1 expression levels, as well as in the ratio of long to short NQO1 isoforms (Fig 2g). Consistent with our previous results, knockdown of NQO1-AS had the opposite effect, decreasing the ratio of long to short isoforms (Fig 2h). Together, these results suggest that HNRNPC binding to NQO1 mRNA favors a truncation of the 3’UTR, and that binding of NQO1-AS prevents this interaction and favors the full-length isoform.

### HNRNPA2B1 stabilizes the long NQO1 isoform and promotes NQO1 expression

Based on our observation that NQO1-AS binding impacts the poly(A) site selection as well as the stability of NQO1 mRNA, we hypothesized that the long and short 3’UTR NQO1 isoforms are differentially regulated, and that the longer 3’UTR isoform is more stable. To assess this, we analyzed CLIP-seq datasets to look for RBP binding sites that are present in the distal portion of the NQO1 3’UTR^28^. This analysis revealed several binding sites for HNRNPA2B1, which we had previously shown to bind and stabilize mRNA in MDA-MB-231 cells^20^ (Fig 3a). Next, we performed an unbiased analysis of the NQO1 3’UTR distal to the region complementary to NQO1-AS using DeepBind^29^ which revealed a strong match to the HNRNPA2B1 consensus motif (Fig S3a). Analyses of the METABRIC and TCGA-BRCA datasets showed a positive correlation between HNRNPA2B1 and NQO1 expression (Fig S3b-c), and, in the case of the TCGA-BRCA dataset, between HNRNPA2B1 expression and NQO1 stability (Fig S3c). These datasets did not show any correlation between HNRNPA2B1 and NQO1-AS expression (Fig S3d). To experimentally test this, we measured NQO1 mRNA stability and expression after HNRNPA2B1 knockdown. Consistent with its role as an mRNA stabilizer, silencing of HNRNPA2B1 increased NQO1 mRNA decay rate and decreased its expression (Fig 3b-c). Notably, silencing of HNRNPA2B1 resulted in a greater decrease in the expression of the full length NQO1 transcript relative to the truncated isoform (Fig 3d). These results support the hypothesis that the two isoforms that result from alternate NQO1 poly(A) site selection are differentially regulated, and that HNRNPA2B1 is responsible for stabilizing the longer isoform.

**Figure 3.**
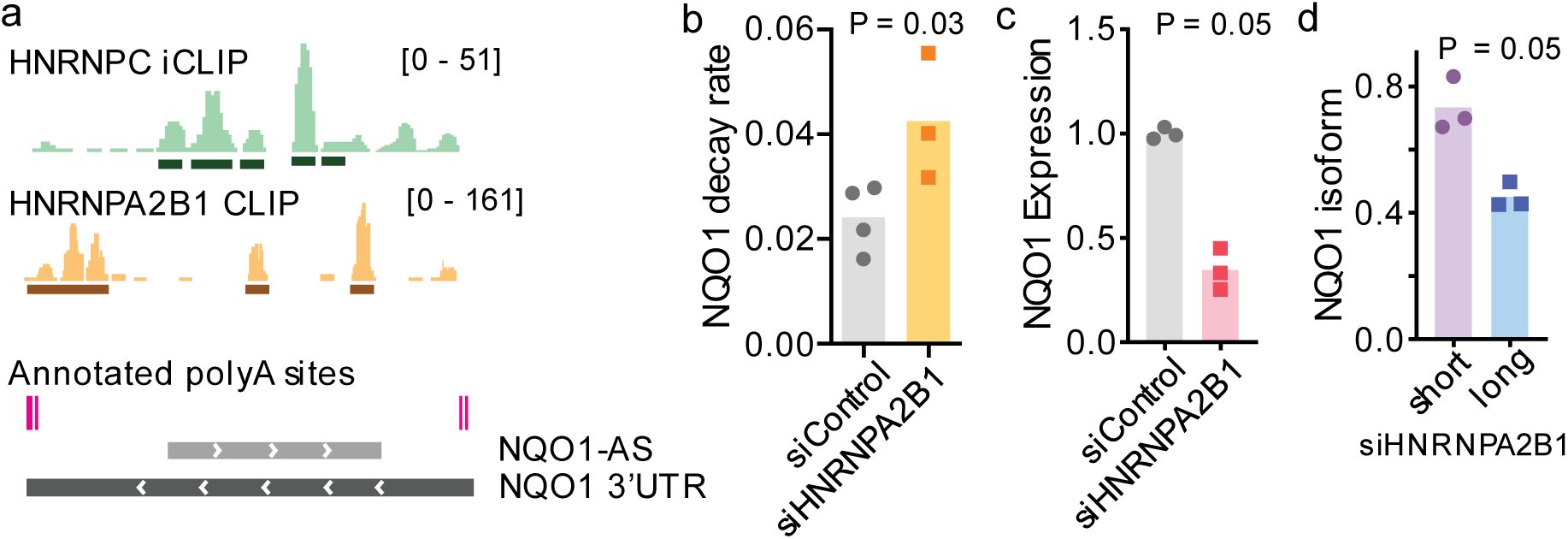
HNRNPA2B1 binds and stabilizes the long NQO1 isoform and increases NQO1 expression. (a) Read distribution in HNRNPC iCLIP and HNRNPA2B1 CLIP experiments relative to annotated poly(A) sites and NQO1-AS complementary region along the NQO1 3’UTR. (b) NQO1 mRNA decay rate as measured by qPCR after α-amanitin treatment of MDA-LM2 cells with and without knockdown of HNRNPA2B1. N = 4. (c) qPCR showing NQO1 expression in MDA-LM2 cells with and without knockdown of HNRNPA2B1. N = 3. (d) qPCR showing NQO1 long and short isoform fractions in HNRNPA2B1 knockdown MDA-LM2 cells. N = 3.

Taken together, our results suggest a new post-transcriptional mechanism for the upregulation of NQO1 in highly metastatic breast cancer cells relative to their poorly metastatic counterparts. By up-regulating NQO1-AS, breast cancer cells inhibit the interaction between HNRNPC and NQO1 mRNA, increasing levels of the full-length NQO1 transcript and its subsequent stabilization by HNRNPA2B1. Thus, expressing NQO1-AS enables cancer cells to decouple NQO1 expression levels from the broader HNRNPC regulon.

### CTCF promotes NQO1-AS transcription in highly metastatic cells

We next sought to identify the factor(s) responsible for NQO1-AS up-regulation in metastatic cells. To do this, we used DeepBind^29^ to perform an unbiased analysis of the putative NQO1-AS promoter, and searched for all known transcription factor binding sites that may act as activators. We discovered a strong match to the CTCF transcription factor consensus motif, and found evidence of CTCF binding to the NQO1-AS promoter in multiple ChIP-seq datasets (Fig 4a; ENCODE). Additionally, we found that CTCF expression was strongly and significantly correlated with NQO1-AS expression in the TCGA breast cancer dataset (Rho=0.4, P=1e-16; Fig 4b). As CTCF is significantly up-regulated in highly metastatic MDA-LM2 cells relative to their parental cells (Fig 4c), we hypothesized that CTCF drives NQO1-AS up-regulation. Further analysis of the TCGA-BRCA and METABRIC datasets revealed a positive correlation between CTCF expression and NQO1 stability and expression (Fig S4a-c), consistent with a model in which CTCF drives the upregulation of NQO1-AS. To test our hypothesis experimentally, we silenced CTCF in MDA-LM2 cells and observed a subsequent reduction in both NQO1-AS and NQO1 expression (Fig 4d-e). We then performed ChIP-qPCR to look for binding of CTCF to the NQO1-AS promoter in MDA-LM2 and MDA-Par cells. Consistent with the pattern of NQO1-AS expression, we saw significantly higher CTCF binding to the region upstream of NQO1-AS in the highly metastatic line (Fig 4f), consistent with CTCF driving NQO1-AS expression.

**Figure 4.**
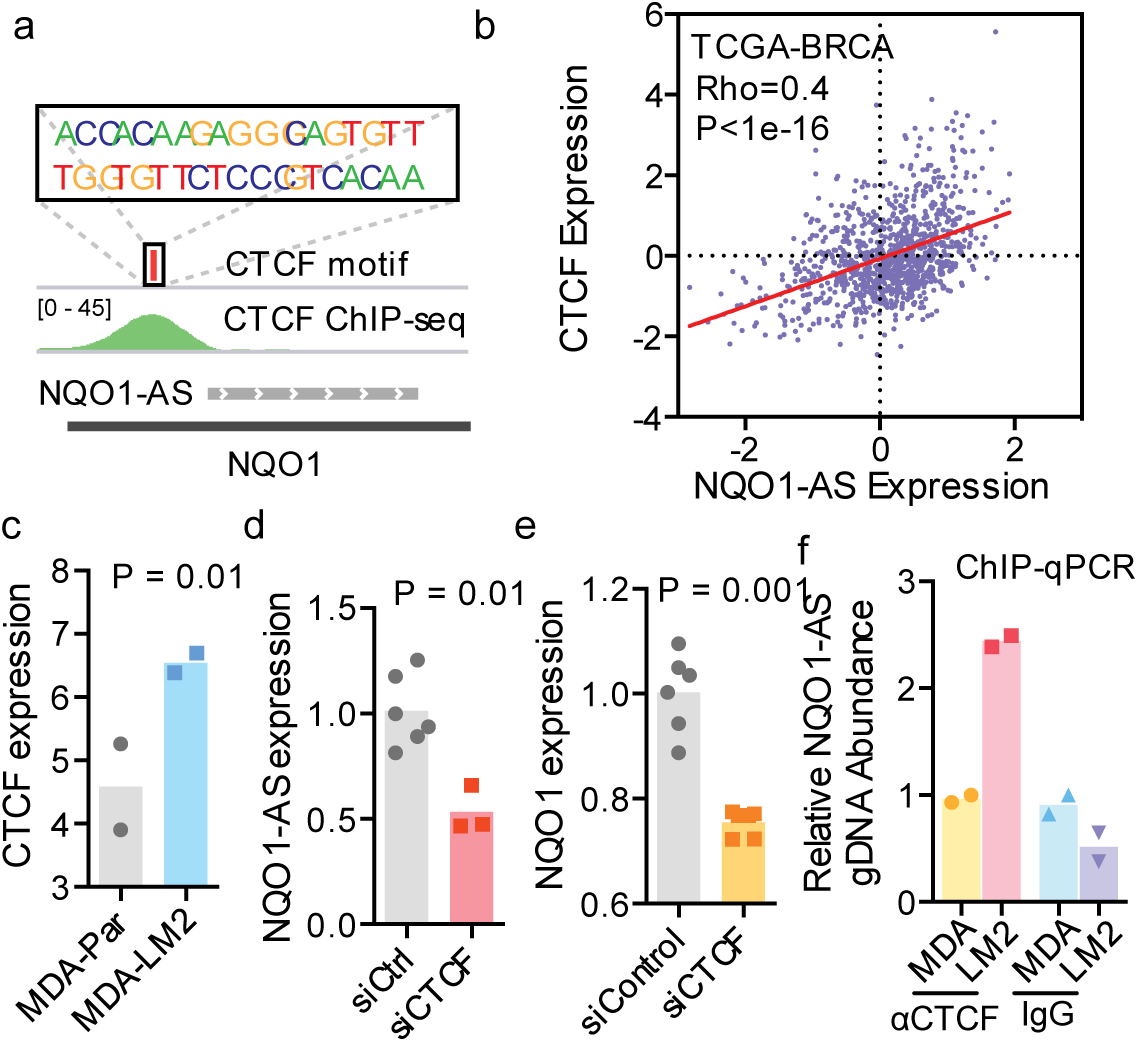
CTCF binding promotes NQO1-AS transcription in highly metastatic cells. (a) CTCF consensus motif relative to CTCF ChIP-seq reads and NQO1-AS complementary region. (b) Correlation between CTCF expression and NQO1-AS expression in TCGA-BRCA dataset. (c) qPCR showing CTCF expression levels in MDA-Par and MDA-LM2 cells. N = 2. (d) qPCR showing NQO1-AS expression in MDA-LM2 cells with and without knockdown of CTCF. N = 6. (e) qPCR showing NQO1 expression in MDA-LM2 cells with and without CTCF knockdown. N = 6. (f) ChIP-qPCR with precipitation of CTCF or IgG (control) in MDA-Par and MDA-LM2 cells. qPCR primers are targeting NQO1-AS promoter region. N = 2.

### NQO1 and NQO1-AS promote metastatic lung colonization

We next investigated the impact of the up-regulation of NQO1 on breast cancer progression. To assess this relationship experimentally, we performed *in vivo* lung colonization assays with NQO1 knockdown and control cells in the MDA-LM2 background. While knocking down NQO1 did not affect the *in vitro* proliferation rate of the cells (Fig S5a), we observed a significant reduction in metastatic burden in the lungs of mice injected with NQO1 knockdown cells (Fig 5a). We repeated this experiment in the HCC1806-LM2 breast cancer cell line to confirm that these results are not cell line-specific, and we again observed that the NQO1 knockdown cells had a significantly reduced capacity for metastatic lung colonization (Fig 5b). Next, we performed lung colonization assays with parental MDA-MB-231 cells overexpressing NQO1. We observed that mice injected with cells overexpressing NQO1 had significantly higher metastatic colonization capacity than control cells (Fig S5b). Given our previous results showing that NQO1-AS stabilizes NQO1 mRNA and leads to increased expression of NQO1, we expected to see similar results in lung colonization assays with NQO1-AS knockdown cells. Consistently, we found that NQO1-AS knockdown in MDA-LM2 cells resulted in decreased lung colonization capacity despite having a similar proliferation rate as control cells *in vitro* (Fig S5c, Fig 5c). Together, our results suggest that increased expression of NQO1 and NQO1-AS promotes the breast cancer metastatic colonization.

**Figure 5.**
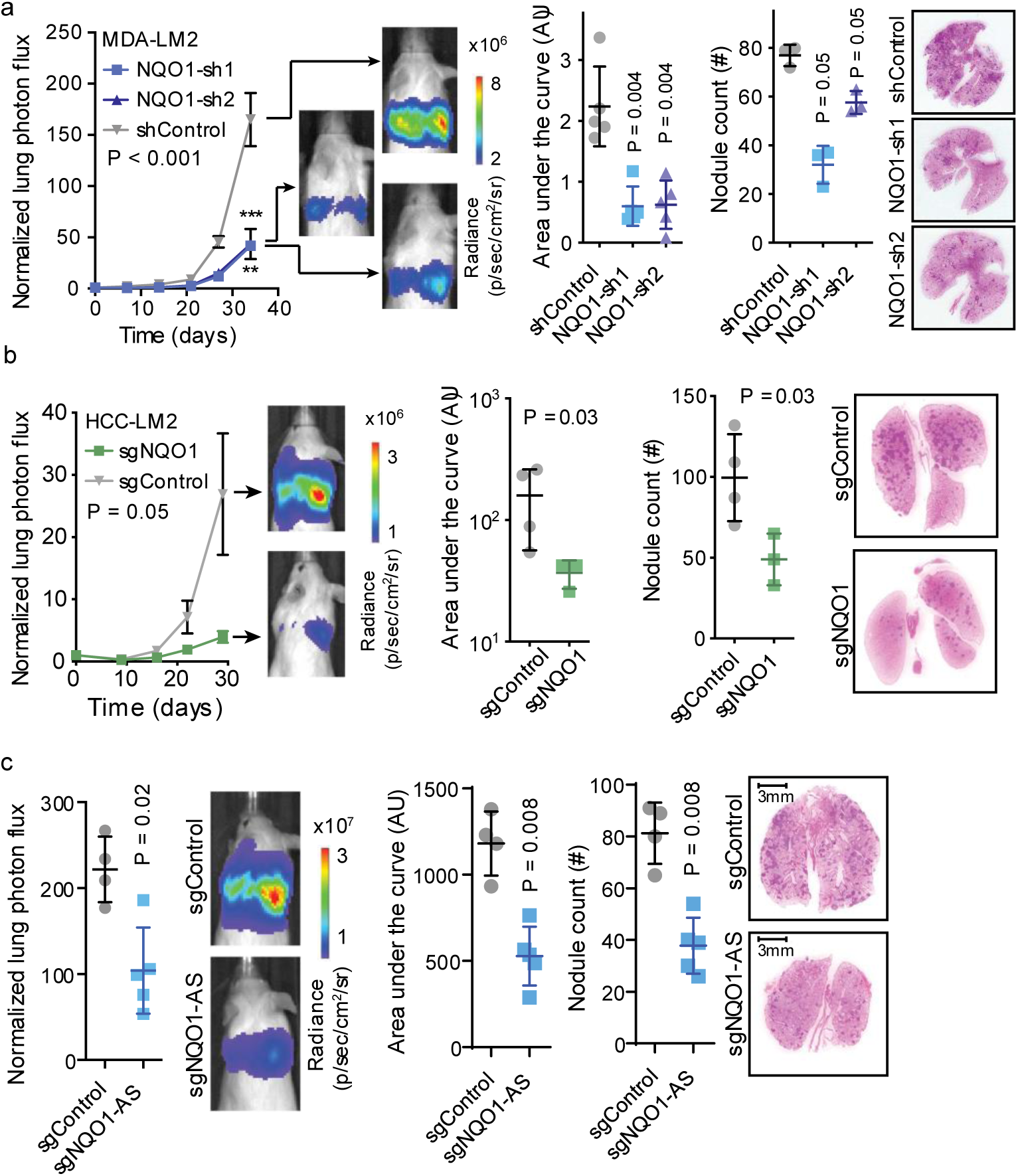
NQO1 and NQO1-AS promote metastatic lung colonization. (a) Lung colonization assays comparing the *in vivo* metastatic colonization capacity of NQO1 knockdown and control MDA-LM2 cells. In vivo measurement of bioluminescence (by cancer cells that constitutively express luciferase) is used as a proxy for tumor burden and validated by endpoint lung histologic sections. N = 5. (b) Lung colonization assays comparing the *in vivo* metastatic capacity of NQO1 knockdown and control HCC1806-LM2 cells. N = 4. (c) Lung colonization assays comparing the *in vivo* metastatic capacity of NQO1-AS knockdown and control MDA-LM2 cells. N = 4.

We next asked whether NQO1 expression plays a role in primary tumor growth *in vivo*. To address this, we injected NQO1 knockdown and control MDA-LM2 cells into mouse mammary fat pads, and measured the tumors over time. We observed no significant difference in growth rate between the two cohorts, indicating that NQO1 expression does not affect primary tumor growth (Fig S5d). This result is unsurprising, since HNRNPC is upregulated during tumorigenesis. Because HNRNPC binds and destabilizes NQO1 mRNA, the upregulation of HNRNPC would be disadvantageous to cancer cells if NQO1 played a significant role in primary tumor growth. Instead, our results indicate that NQO1 is required primarily for metastasis, and that its decoupling from the HNRNPC regulon via upregulation of NQO1-AS is a contributor to metastatic transformation.

### NQO1-mediated metabolic remodeling protects cancer cells from ferroptosis

Having observed that NQO1 promotes breast cancer metastasis, we next sought to determine the mechanism by which it exerts its pro-metastatic effects. NQO1 is a chemoprotective enzyme involved in cellular defense against oxidizing agents^30^. In addition to reducing a wide range of substrates and counteracting the production of reactive oxygen species (ROS), it also helps scavenge superoxides^31^. NQO1 is directly involved in NADPH metabolism, and its up-regulation in highly metastatic cells may therefore lead to increased survival under oxidative stress. It was recently shown that melanoma cells reversibly increase their expression of NADPH generating enzymes to boost their capacity to withstand oxidative stress^32^. Given that breast cancer cells also experience oxidative stress during metastasis, both in circulation and during lung colonization^33^, it is possible that NQO1 plays a similar role in breast cancer metastasis. We investigated this possibility using NQO1 knockdown and control cells in both MDA-LM2 and HCC1806-LM2 backgrounds. As mentioned above, silencing NQO1 did not impact cell proliferation rates in culture (Fig S5a), but we observed a significantly higher level of total reactive oxygen species in NQO1 knockdown cells relative to control cells as measured by a fluorometric assay (Fig 6a-b). Additionally, we observed a significant decrease in the survival of NQO1 knockdown cells when exposed to oxidative stress induced by hydrogen peroxide treatment (Fig S6a). MDA-Par cells, which endogenously express NQO1 at a lower level compared to MDA-LM2 cells, also showed a lower tolerance to H_2_O_2_ treatment (Fig S6a). We repeated these experiments in NQO1-AS knockdown MDA-LM2 cells, and, consistent with the dependence of NQO1 levels on NQO1-AS, found that NQO1-AS knockdown cells had higher baseline ROS levels and increased sensitivity to H_2_O_2_ (Fig 6a, Fig S6b).

**Figure 6.**
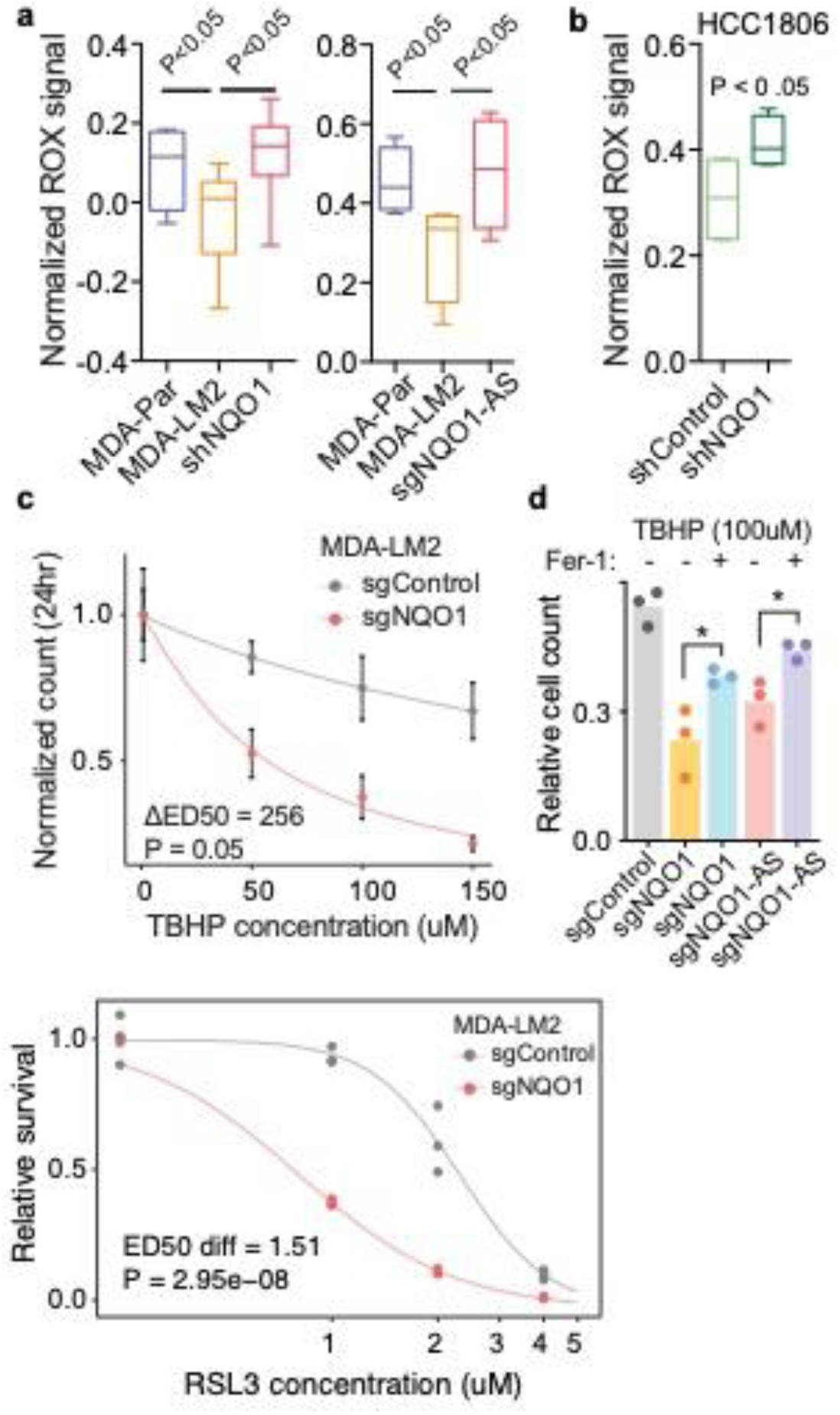
NQO1 protects cancer cells from ferroptosis. (a) Reactive oxygen species as measured by Invitrogen’s CellRox assay in A-Par, MDA-LM2, MDA-LM2 NQO1 knockdown, and MDA-LM2 NQO1-AS knockdown cells. N = 4. (b) Reactive oxygen species as measured by a fluorometric (CellRox, Invitrogen) in HCC1806-LM2 NQO1 knockdown and control cells. (c) Graph showing sensitivity of NQO1 knockdown and control MDA-LM2 cells to TBHP. N = 6. (d) Graph showing impact of Fer-1 pre-treatment on TBHP sensitivity of NQO1 and NQO1-AS knockdown and control MDA-LM2 cells. N = 6. (e) Graph showing sensitivity of NQO1 knockdown and control MDA-LM2 cells to RSL3. N = 6.

Ferroptosis, a non-apoptotic form of programmed cell death, has recently been shown to be a consequence of excessive oxidative stress in metastatic breast cancer^34,35^. Given the role of NQO1 in protecting breast cancer cells from ROS, we hypothesized that NQO1 may be promoting lung metastasis by preventing ferroptotic cell death. To test this, we treated NQO1 knockdown and control cells with tert-butyl hydroperoxide (TBHP), a known inducer of oxidative stress and ferroptosis^36^. In both MDA-LM2 and HCC1806-LM2 backgrounds, NQO1 knockdown cells were significantly more sensitive to TBHP compared to control cells (Fig 6c, Fig S6c). When we pre-treated NQO1 and NQO1-AS knockdown cells with Ferrostatin-1 (Fer-1), a known inhibitor of ferroptosis^37^, we saw a decrease in sensitivity to TBHP treatment, indicating that a portion of the cell death we observed was likely due to ferroptosis (Fig 6d). Liproxstatin-1, another inhibitor of ferroptosis^37^, similarly blunted the sensitivity of MDA-LM2 cells to TBHP (Fig S6e). To address the possibility that TBHP induced apoptosis in the treated cells, we assayed the caspase activities of treated and untreated MDA-LM2 NQO1 knockdown and control cells using a luminescence-based assay. While the TBHP treated cells showed increased caspase activity, there was no difference between the NQO1 knockdown and control cells, indicating that NQO1 selectively protects against ferroptosis (Fig S6g). Next, we tested the TBHP sensitivity of MDA-MB-231 cells that have been in vivo selected to metastasize to the bone (BoM) and brain (BrM2)^38^ and compared them to the MDA-Par and MDA-LM2 lines. We found that there was no significant difference in the sensitivities of MDA-Par, MDA-BoM, and MDA-BrM2 to TBHP, and that only the MDA-LM2 line showed a decreased sensitivity (Fig S6d). As the MDA-LM2 cell line has been *in vivo* selected to metastasize to the lung, this result suggests that breast cancer metastases to the lung are uniquely resistant to ferroptosis, consistent with the high oxidative stress that exists in the lung environment. To provide additional evidence that NQO1 protects cells from ferroptosis, we tested the sensitivity of NQO1 knockdown and control MDA-LM2 cells to two established inducers of ferroptosis: RSL3 (Fig 6e) and Erastin (Fig S6f)^39^. Consistent with our previous results, knockdown of NQO1 significantly increased the cells’ sensitivity to both compounds, indicating that NQO1 knockdown cells are more vulnerable to ferroptotic cell death.

Because ferroptosis is associated with a disruption of normal cellular metabolism, we performed LC/MS-based metabolic profiling of breast cancer cells^40^. Consistent with the role of NQO1 as a regulator of the cell’s redox state, we observed a significant change in several redox-dependent metabolites upon knockdown of NQO1 (Fig 7a, S7b). We found that, in both the MDA-LM2 and HCC1806-LM2 backgrounds, NQO1 knockdown cells exhibited significantly higher NADPH, malate, and hydroxyproline levels relative to controls (Fig 7a). NQO1 regulates the redox state of the cell primarily by coupling the reduction of ROS to the oxidation of NADPH, and an increase in NADPH upon knockdown of NQO1 (Fig S7b) is consistent with this role. To our knowledge, NQO1 is not directly involved in the metabolism of malate or hydroxyproline, but it is possible that the buildup of these metabolites in NQO1 knockdown cells is due to a disruption in the metabolism of ubiquinone (CoQ_10_). One of the mechanisms by which NQO1 protects against oxidative damage is through the reduction of CoQ_10_ to ubiquinol (QH_2_), which subsequently acts as an antioxidant^41^. CoQ_10_ is also used as an electron carrier by proline dehydrogenase 2 (PRODH2), which catalyzes the first step of hydroxyproline catabolism^42^, and by the electron transport chain (ETC), of which malate dehydrogenase is an essential component. Decreased turnover of CoQ_10_ in the absence of NQO1 activity may therefore perturb these metabolic systems and cause the observed increase in hydroxyproline and malate. To explore whether NQO1’s impact on ferroptosis relates to CoQ_10_ metabolism, we pretreated NQO1 knockdown and control MDA-LM2 cells with either CoQ_10_ or QH_2_ and measured their sensitivity to TBHP. We found that pretreatment with QH_2_ rescued the increased sensitivity to TBHP that we had previously observed in NQO1 knockdown cells relative to controls (Fig 7b). Pretreatment with CoQ_10_, however, resulted in a mild but statistically insignificant rescue of this phenotype. These results suggest that decreased NQO1 activity limits a cell’s ability to utilize CoQ_10_ as an antioxidant, and that NQO1 protects against ferroptosis at least in part through the production of QH_2_.

**Figure 7.**
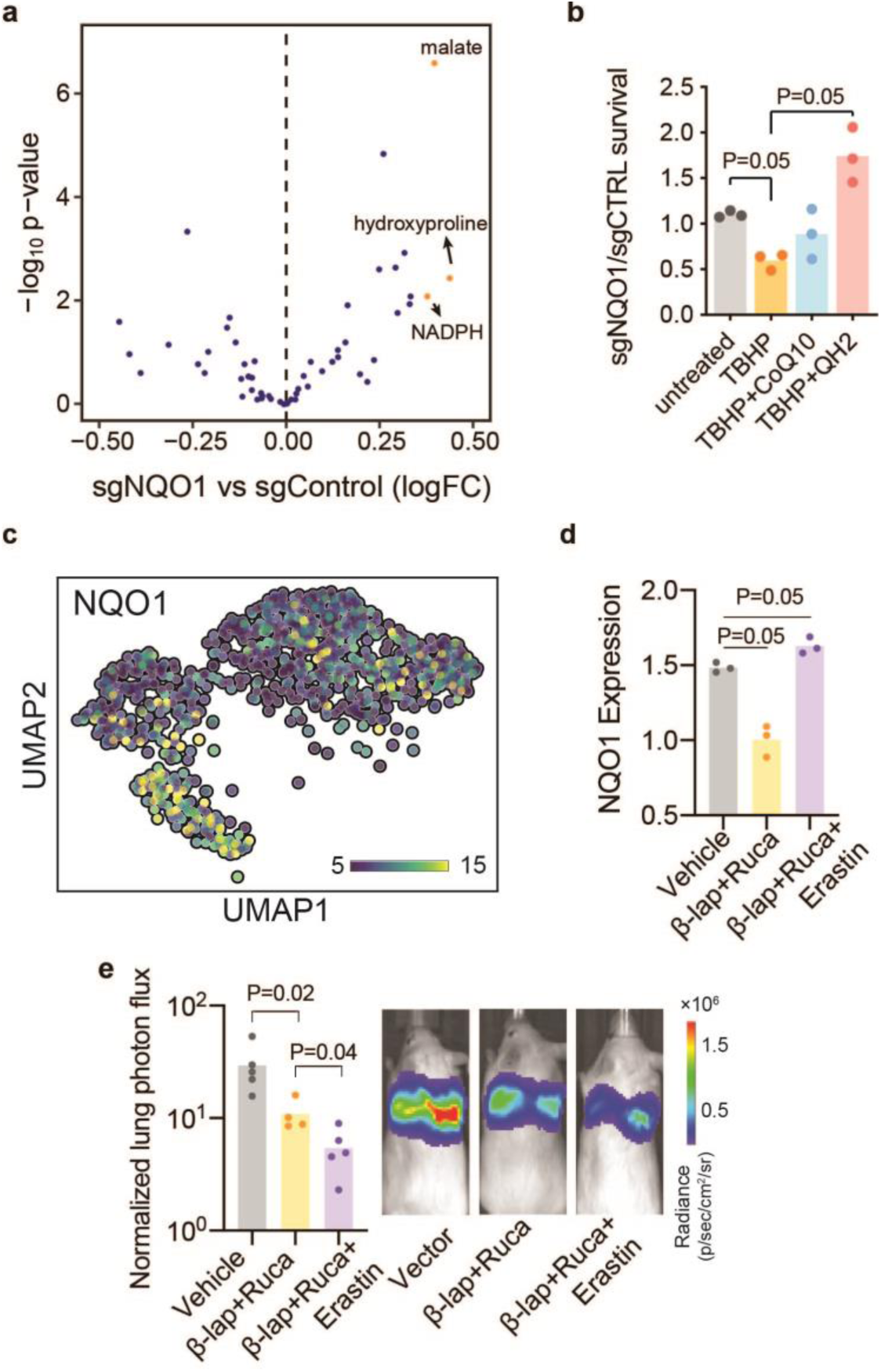
NQO1 mediates metabolic remodeling in breast cancer cells. (a) Volcano plot showing change in metabolic landscape of MDA-LM2 and HCC1806-LM2 cells in response to NQO1 knockdown. Metabolite levels were measured using LC/MS-based metabolic profiling. N = 3. (b) Ratio of surviving NQO1 knockdown to control MDA-LM2 cells after pretreatment for 24h with 10 µM ubiquinone (CoQ_10_) or ubiquinol (QH_2_), followed by treatment for 24h with 150µM TBHP. N = 3. (c) UMAP plot generated from single-cell sequencing data in MDA-Par cells. Cells are colored according to relative NQO1 expression. (d) qPCR showing NQO1 expression level of MDA-LM2 cells after treatment with β-Lapachone + Rucaparib, β-Lapachone + Rucaparib + Erastin, or vector control. N = 3. (e) Luciferase fluorescence signal from *in vivo* lung colonization assay with MDA-Par cells and subsequent treatment with β-Lapachone + Rucaparib, β-Lapachone + Rucaparib + Erastin, or vehicle control. Representative images from each cohort are shown. N = 5.

Next, we treated cells from both backgrounds with TBHP and repeated the metabolic profiling. As expected, in the TBHP-treated cells there were changes in redox-dependent metabolites, and these changes were relatively consistent across the two backgrounds (Fig S7a). This result highlights the extent of the metabolic remodeling that occurs with changes in NQO1 expression. Breast cancer cells that are able to upregulate NQO1 alter their metabolome to favor the reduction of reactive oxygen species and protection against ferroptosis.

### The role of NQO1 as a protective agent against ferroptosis can be exploited therapeutically

Given the role of NQO1 in promoting breast cancer metastasis, we explored ways in which it could be targeted therapeutically. Through its enzymatic activity, NQO1 can convert certain compounds to a more potent form within cancer cells. For example, β-Lapachone is metabolized by NQO1 into an unstable hydroquinone that spontaneously oxidizes and generates superoxide^43^. Through this futile cycle, β-Lapachone generates ROS and depletes NADPH in cancer cells, which in turn leads to programmed necrosis^44^. This compound has proved effective in a number of cancers with increased NQO1 expression; however, its sustained use at high concentrations causes anemia in both humans and animal models^45^. Recently, it has been shown that combining β-Lapachone with poly(ADP-ribose) polymerase (PARP) inhibitors results in synergistic anti-tumor activity, as DNA lesions resulting from ROS cannot be repaired^46^. Given our results showing that NQO1 is protective against ferroptosis, we hypothesized that low levels of Erastin, a ferroptosis inducing agent, could enhance the potency of the combined β-Lapachone/PARP inhibitor treatment. Additionally, single-cell RNA sequencing revealed a high degree of heterogeneity in NQO1 expression in MDA-MB-231 cells (Fig 7c). Cells expressing high levels of NQO1 clustered together in a UMAP analysis, and they overlapped with the cluster formed by MDA-LM2 cells when these parental and highly metastatic lines were analyzed together (Fig S7c). This result suggests that the subpopulation of MDA-MB-231 cells that express high levels of NQO1 are the precursors to the MDA-LM2 derivatives, and they are therefore an important subset to target therapeutically. Targeting PARP and NQO1 while inducing ferroptosis could therefore enable the simultaneous targeting of subpopulations of cancer cells with high and low NQO1 expression. To test this, we first treated MDA-LM2 cells with either β-Lapachone + Rucaparib (a PARP inhibitor), Erastin, or vehicle control, and then measured the relative NQO1 expression level of the surviving cells by qPCR (Fig 7d). We found that the cells that survived β-Lapachone + Rucaparib treatment had a relatively low NQO1 level, indicating that the regimen had killed the population of cells that were upregulating NQO1. Conversely, the cells that survived Erastin treatment had a relatively high NQO1 level, indicating that Erastin had killed the population of cells under-expressing NQO1. This result suggests that β-Lapachone and Erastin target at least partially distinct subpopulations, and may therefore have synergistic activity if combined. To test this possibility, we performed a lung colonization assay with MDA-Par cells in three cohorts of mice: one receiving β-Lapachone + Rucaparib; one receiving β-Lapachone + Rucaparib + Erastin; and one control. As we hypothesized, the mice receiving β-Lapachone, Rucaparib, and Erastin in combination had a significantly lower metastatic burden than the other cohorts, as measured both by *in vivo* bioluminescence measurements (Fig 7e) and endpoint histologic staining (Fig S7d). This result suggests that the role of NQO1 as a protective agent against ferroptosis can be exploited in combination with established therapies.

### Increased NQO1-AS and NQO1 expression is associated with metastasis in clinical samples

To assess the clinical relevance of NQO1 in breast cancer metastasis we examined the relationship between NQO1 expression and cancer progression in clinical samples. We analyzed RNA-seq data from four patient derived xenografts (PDX) lines^47^, two of which were poorly metastatic (HCI-002 and HCI-004), and two of which were highly metastatic (HCI-001 and HCI-010), and found that the highly metastatic PDXs expressed significantly higher levels of NQO1 and NQO1-AS (Fig S8a). Our analysis of the TCGA-BRCA dataset showed significantly lower survival for patients with tumors expressing high levels of either NQO1-AS or NQO1 (Fig 8a-b). Similarly, both NQO1-AS and NQO1 expression were positively correlated with disease stage (Fig 8c). Our analysis of the METABRIC dataset yielded similar results, with high NQO1 expression associated with lower disease-free survival and higher disease stage (Fig 8d-e). A negative association between NQO1 expression and survival was also observed in multiple other breast cancer gene expression datasets (Fig S8b-c). We also performed qPCR to assay NQO1 levels in a panel of cDNA derived from breast cancer tissue (Origene). We found that NQO1 expression was associated with disease stage, and that samples from higher disease stages were significantly more likely to express detectable levels NQO1-AS (Fig 8f-g). We performed immunohistochemistry to assess NQO1 expression in breast cancer tissue from progressive cancer stages. Consistent with our earlier analyses, we found that there was significantly higher NQO1 expression in lymph node metastases, invasive lobular carcinoma, and invasive ductal carcinoma than in ductal carcinoma in situ or non-neoplastic breast tissue (Fig S8c). Together, these results indicate that our findings in established cell lines are reproducible in clinical samples.

**Figure 8.**
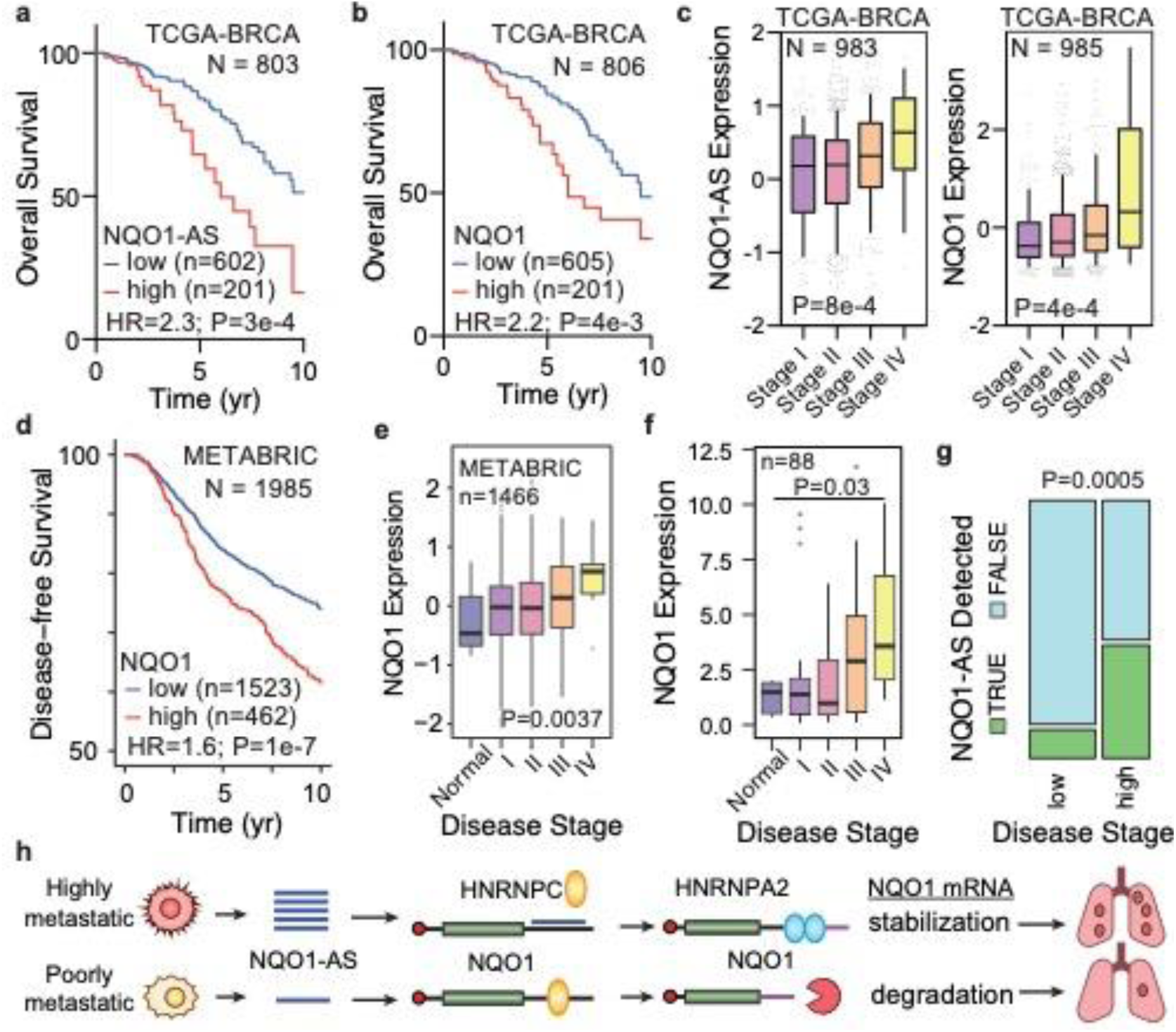
NQO1-AS and NQO1 expression are associated with metastasis in clinical samples. (a) Kaplan-Meier curve showing survival rates of patients with tumors expressing high or low levels of NQO1-AS. Data from TCGA-BRCA. N = 803. (b) Kaplan-Meier curve showing survival rates of patients with tumors expressing high or low levels of NQO1. Data from TCGA-BRCA. N = 806. (c) NQO1-AS and NQO1 expression in tumors associated with stage I, II, III, and IV breast cancer. Data from TCGA-BRCA. N = 983. (d) Kaplan-Meier curve showing disease-free survival of patients with tumors expressing high or low levels of NQO1. Data from METABRIC. N = 1985. (e) NQO1 expression in normal cells and cells associated with stage I, II, III, and IV breast cancer. Data from METABRIC. N = 1466. (f) NQO1 expression in normal cells and cells associated with stage I, II, III, and IV breast cancer. Data from BCRT102 and BCRT103. N = 88. (g) Graph showing fraction of tumors with detectable NQO1-AS in low and high stage breast cancer. Data from Origene Tissue Scan (BCRT102 and BCRT103). N = 88. (h) Schematic showing the mechanism by which upregulation of NQO1-AS promotes breast cancer metastasis.

## Discussion

We have developed a computational method to identify expressed antisense RNAs, and have applied it to models of breast cancer metastasis. Here, we have focused our attention on one of these identified antisense RNAs, NQO1-AS, which is both cancer-specific and associated with breast cancer progression. We hypothesized that, since NQO1 is similarly associated with breast cancer progression, it may be upregulated in highly metastatic cells through a pathway involving a direct interaction with NQO1-AS. Taken together, our data supports a model in which NQO1-AS binds to the 3’ UTR of NQO1, preventing the binding of HNRNPC, thereby favoring use of the distal polyadenylation site and increased abundance of the longer NQO1 isoform. HNRNPA2 1 can then bind to the long 3’UTR, stabilizing the mRNA and increasing the level of NQO1 protein. Breast cancer cells exploit this pathway during metastatic progression, enabling them to tolerate higher levels of oxidative stress. Cells that fail to upregulate NQO1 in this way undergo ferroptosis, and those that successfully colonize the lungs become dependent on this pathway for survival. As this dependence evolves, metastatic cancer cells are vulnerable to drugs that target this pathway. In our preliminary experiments, we have shown that combining NQO1 and PARP inhibitors with a ferroptosis inducing agent can significantly reduce the metastatic capacity of breast cancer cells, and this combination could form the basis for a potent therapeutic regimen.

Antisense RNAs are emerging as an important class of regulatory molecules with a wide variety of functional roles, and, although more work is needed to fully understand how they contribute to cancer progression, it has become clear that they can alter gene expression at every level^48^. The pathway that we have identified here represents a novel mechanism by which cancer cells exploit antisense RNAs to regulate gene expression post-transcriptionally. Silencing NQO1-AS in our model of breast cancer metastasis has a profound effect on the metastatic capacity of cancer cells, and the magnitude of this impact highlights the significance of antisense RNA-mediated gene expression changes. In this case, upregulation of an antisense RNA enables cancer cells to disrupt healthy regulatory pathways and undergo a metabolic remodeling that makes them better suited to invade distal organs. Although we have focused here on a single pathway, the pool of antisense RNAs that we have identified likely includes a number of molecules that play a role in regulating gene expression through direct RNA-RNA interactions. Further investigation of these interactions may lead to a better understanding of how cells transition from healthy to diseased states, and may reveal new targets for the development of improved therapies.

In addition to gene expression dysregulation, metabolic reprogramming is an essential step in tumorigenesis and cancer progression^49^. As cancer cells alter their metabolic landscape, they become resistant to some stressors and sensitive to others. Since its discovery, ferroptosis has drawn considerable interest as a form of regulated cell death that can be induced in multiple treatment resistant cancers^50^. Triple negative breast cancer, which is characteristically resistant to targeted therapies, has been shown to undergo ferroptosis in response to GPX4 inhibition with Erastin^51^. Here, we have shown that MDA-MB-231 cells are able to protect themselves from ferroptotic cell death by overexpressing NQO1 and remodeling their metabolic profile. In so doing, they become increasingly metastatic and are better able to colonize the lungs, but, importantly, this process makes the metastatic cells dependent upon NQO1 upregulation. This dependence creates an attractive therapeutic opportunity, as NQO1 can be targeted with β-Lapachone, “unmasking” the cancer cells’ sensitivity to rastin. In a mouse model, we have shown that adding rastin to a therapeutic regimen of β-Lapachone and Rucaparib results in a significantly lower tumor burden. The use of β-Lapachone in humans has been limited by toxicity when taken at high doses for a sustained period, but its inclusion in a more potent combination of drugs may enable the dose to be lowered to a level that is tolerable to patients. More work will be required to identify the optimal drug combination, but we have demonstrated the viability of this therapeutic strategy.

## Acknowledgements

We acknowledge the UCSF Center for Advanced Technology (CAT) and the Chan Zuckerberg Biohub for high throughput sequencing and other genomic analyses. We thank Byron Hann and the Preclinical Therapeutics core as well as the Laboratory Animal Resource Center (LARC) at UCSF. We are also grateful for the genomic data contributed by the TCGA Research Network, including donors and researchers. We acknowledge support from our colleagues at the Helen Diller Family Comprehensive Cancer Center. We also acknowledge Jonathan Weissman and Luke Gilbert for CRISPRi constructs. We acknowledge Dr. Brent R. Stockwell for his input on studying ferroptosis. We also acknowledge the Ludwig Cancer Research Institute. This work was supported by grants from the NIH (R00CA194077 and R01CA24098) and ACS (130920-RSG-17-114-01-RMC) to H.G., as well as grant R01CA163591 from the NIH to J.D.R.

