## Supplementary Information for "A sense-antisense RNA interaction promotes breast cancer metastasis via regulation of NQO1 expression"

### Methods

#### *Tissue culture*

All cells were cultured at 37 °C in a humidified incubator with 5% $CO_2$. MDA-MB-231, MDA-LM2, CN34, CN34-LM1a, and 293LTV cell lines were grown in DMEM supplemented with 10% FBS, penicillin (100 units/mL), streptomycin (100 µg/mL) and amphotericin (1 µg/mL). HCC1806 and HCC1806-LM2 cell lines were grown in RPMI-1640 supplemented with 10% FBS, L-glutamine (2mM), sodium pyruvate (1mM), penicillin (100 units/mL), streptomycin (100 µg/mL) and amphotericin (1 µg/mL).

#### *Stable and transfected cell lines*

Cells were transduced using the ViraSafe lentiviral packaging system (Cell Biolabs). NQO1 was silenced both with shRNA and with CRISPRi constructs expressing the sequences in Table 2. NQO1-AS and HNRNPC were similarly silenced by expressing the guide sequences from the Table 2 in CRISPRi constructs. CTCF and HNRNPA2B1 were silenced using siRNA. NQO1 was overexpressed using a CRISPR-activation construct. NQO1-AS overexpression was accomplished by cloning the sequence from the attached file into the PLX302 backbone.

#### *Quantitative RT-PCR*

Transcript levels were measured using quantitative RT-PCR by reverse transcribing total RNA into cDNA (SuperScript III or Maxima H Minus, Invitrogen), then using Perfecta SYBR green supermix (QuantaBio) for amplification, per the manufacturer’s instructions. HPRT1 was used as the endogenous control. All primer sequences in Table 1.

#### *RNA-seq Library Preparation*

RNA-seq libraries were prepared using the ScriptSeq-v2 kit (Illumina) using RNA that had been rRNA depleted using the Ribo-Zero Gold kit (Illumina). Libraries were sequenced on an Illumina HiSeq4000 instrument at the UCSF Center for Advanced Technologies.

#### *3’end-seq*

RNA-seq libraries were constructed with QuantSeq Rev Kit (Lexogen) following the manufacturer’s protocol and sequenced on Illumina HiSeq sequencer in the Center for Advanced Technology (UCSF). Cutadapt was used to remove short and low-quality reads, that were then aligned to hg38 reference genome using salmon and compared using DESeq2. Reads mapping to the annotated NQO1 polyA sites were extracted and compared between conditions using logistic regression.

#### *Single-cell RNA seq*

MDA-Par cells were split into biological replicates, barcoded, and scRNA-seq libraries were prepared with Chromium Next GEM Single Cell 3’ Kit v3 (10x Genomics). The libraries were sequenced on an Illumina NovaSeq sequencer in the Chan Zuckerberg Biohub.

***Single-cell RNA seq analysis***

*Raw Data Processing*
CellRanger v3.0 (10x Genomics) was utilized for cell barcode filtering, read alignment, UMI counting, and generating a digital gene expression matrix from raw fastq files. Reads were aligned to the human reference genome hg38, using CellRanger-provided annotations for gene features. Reads were assigned to cells based on their cell barcodes, and barcodes that did not appear in the 10x Genomics 3M barcode allow-list were removed.

*Barcode Demultiplexing and Assignment*
Cells were assigned to cell-lines of origin by quantifying the relative proportion of detected genetic barcodes. Unique molecular counts for each barcode were determined from barcode-containing reads, a gaussian kernel density estimation was fit to the frequency of each barcode across all cells. The inter-peak minima of the resulting bimodal distributions were set as the minimum threshold for barcode assignment. Cells were assigned a barcode if the frequency of that barcode exceeded its associated threshold and was 10-fold more frequent than the second most frequently occurring barcode. Cells assigned multiple barcodes were designated as doublets and removed.

*Single Cell Data Pre-Processing and Visualization*
Scanpy[1] was utilized for all preprocessing. Cells expressing fewer than 200 genes or greater than 6000 genes, or with aggregate mitochondrial gene expression was greater than 9% of overall cell expression were removed. Genes expressed in fewer than 3 cells were also removed. Gene expression counts were normalized to 100,000 counts per cell, log-transformed after adding a pseudocount, and scaled across cells to unit variance and zero mean. The top 3000 highly variable genes were determined, and cells projected to a lower dimensional representation via principal component analysis with this reduced feature set. UMAP [13], implemented in scanpy.tl.umap with standard parameters, was then applied to cells representedby the minimum number of principal components required to explain the observed variance. Clusters were generated using the Louvain algorithm as implemented in scanpy.tl.louvain.

***Global nuclear run-on assays (GRO-seq)***
Gro-seq was performed as described in Core et al 2008 and Wang et al 2011, with some adaptations: For each sample, nuclei from $1x10^7$ MDA-parental or MDA-LM2 cells were used. All steps were done on ice. Cells were harvested by scraping and then resuspended in swelling buffer (10mM Tris-HCl pH 7.5, 2mM $MgCl_2$, 3mM $CaCl_2$) for 5 minutes on ice, then spun at 500 x g 5 minutes at 4°C. Swelling buffer was aspirated, and 10mL lysis buffer (10mM Tris-HCl pH 7.5, 2mM $MgCl_2$, 3mM $CaCl_2$, 0.5% IGEPAL CA-630, 10% glycerol) was added. The nuclei were then pelleted at 1000 x g for 5 minutes at 4°C. Lysis buffer was aspirated and nuclei resuspended in 1mL freezing buffer (50mM Tris-HCl pH 8.3, 40% glycerol, 5mM $MgCl_2$, 0.1mM EDTA). Nuclei were then pelleted at 1000 x g 5 minutes 4°C. Freezing buffer was aspirated and nuclei resuspended in 100ul of freezing buffer and stored at -80°C. For the run-on assay, the frozen nuclei were thawed on ice and mixed with 100ul reaction buffer (10mM Tris-HCl pH 8.0, 5mM $MgCl_2$, 1mM DTT, 300mM KCl, 0.2units/ul SuperaseIN, 1% sarkosyl, 500uM each ATP, GTP, and Br-UTP, 2uM CTP), then incubated at 30°C for 5 minutes. Reaction was stopped by adding 600ul Trizol LS (Invitrogen), and RNA was isolated per the manufacturer’s protocol. The RNA pellet was resuspended in 20ul H 2 O, and the RNA fractionated by adding 5ul 1M NaOH and incubating on ice for 40 minutes. Reaction was stopped by adding 25ul 1M Tris-HCl pH 6.8. This solution was run through micro biospin P-30 columns (BioRad), then DNase-treated by adding 10ul RQ1 DNase (Promega), 6.7ul 10X DNase reaction buffer, 1ul SuperaseIN and

incubating at 37°C 10 minutes. This was run through a micro biospin P-30 column (BioRad). The RNA was dephosphorylated by adding 5ul Antarctic phosphatase (NEB), 8.5ul 10X phosphatase buffer, 1ul superaseIN, and incubated at 37°C for 1 hour. Labeled RNA was isolated with anti-BrdU agarose beads (Santa Cruz Biotech) that were prepared by incubating in blocking buffer (0.5X SSPE, 1mM EDTA, 0.05% tween-20, 0.1% PVP, 1mg/mL BSA) for 1 hour at 4°C, then resuspended in 500ul binding buffer (0.5X SSPE, 1mM EDTA, 0.05% tween-20). The RNA was added to the prepared beads after the RNA was heated at 65°C for 5 minutes, then placed on ice. The RNA was incubated with the beads with end-over-end rotation for 1 hour at 4°C. The beads were then washed 1X with low salt buffer (0.2X SSPE, 1mM EDTA, 0.05% tween-20), 2X with high salt buffer (0.5X SSPE, 1mM EDTA, 0.05% tween-20, 150mM NaCl), and 2X TET buffer (TE pH 7.4, 0.05% tween-20). RNA was eluted by adding 125ul elution buffer (5mM Tris-HCl pH 7.5, 300mM NaCl, 1mM EDTA, 0.1% SDS, 20mM DTT) and rotating end-over-end at room temp. Elution by repeated for a total of four times. RNA was isolated from the eluate by acid phenol chloroform extraction and precipitation. The RNA pellet was resuspended in 45ul H2O, and then phosphorylated by adding 5.2ul T4 PNK buffer, 1ul superaseIN, 1ul T4 PNK (NEB) and incubating at 37°C for 1 hour. RNA was isolated from the reaction by acid phenol chloroform extraction and precipitation. The RNA was polyA tailed by adding 0.8ul 10X polyA polymerase buffer, 1ul 1mM ATP, 0.5ul SuperaseIN, 0.75ul polyA polymerase (NEB), and incubating at 37°C for 30 minutes. RNA was then reverse transcribed by first adding 1ul 10mM dNTPs, 2.5ul 12.5uM oNTI223 primer, incubating at 75°C 3 minutes, then on ice 1 minute, then adding 2ul 10X superscript III buffer, 2ul 25mM MgCl2, 3ul 0.1M DTT, 0.5ul superaseIN, 1ul superscript III (Invitrogen), and incubating at 48°C for 20 minutes. Excess primer was digested by adding 4ul exonuclease I and incubating at 37°C for 1 hour. RNA was hydrolyzed by adding 1.8ul 1M NaOH and incubating at 98°C for 20 minutes, then adding 1.8ul 1M HCl to neutralize. The resulting cDNA was run on a 10% polyacrylamide TBE-urea gel (Invitrogen) and the region from 110-405nt was excised and the DNA recovered by the crush soak method. The precipitated DNA was resuspended in 7.5ul H2O and circularized by adding 1ul CircLigase buffer, 0.5ul 1mM ATP, 0.5mM MnCl2, 0.5ul CircLigase (Lucigen) and incubated at 60°C for 1 hour, then 80°C for 20 minutes. Linearization was then performed by adding 3.8ul of 100mM KCl, 2mm DTT and 1.5ul APE1 (NEB), and then incubating at 37°C for 1 hour. The cDNA was run on a 10% polyacrylamide TBE-urea gel (Invitrogen) and the region from 125-305nt was excised and the DNA recovered by the crush soak method. The precipitated cDNA was resuspended in 20ul H2O. PCR was then performed by combining 10ul of the cDNA with 4ul 5X phusion HF buffer, 0.4ul 10mM dNTPs, 2ul each 5uM oNTI200 and oNT1201, 1.4ul H2O, 0.2ul phusion HF DNA polymerase (NEB) and running the following cycle: 98°C 30s, then repeat X13 98°C 10s, 60°C 15s, 72°C 15s. PCR product was run on an 8% polyacrylamide TBE gel, and the region from 150-230bp was excised. DNA was extracted by the crush soak method, and the library was sequenced on an Illumina HiSeq2000 with the Illumina small RNA sequencing primer.

#### *Psoralen crosslinking followed by nuclease digestion and RNA ligation*

MDA-Par and MDA-LM2 cells were resuspended in 4 mL ice-cold aminomethyltrioxsalen solution (0.5 mg/mL in PBS) and incubated on ice for 15 minutes in the dark. The mixture was then transferred to a 10 cm prechilled tissue culture plate and irradiated 400mJ/cm$^2$ 254nm UV for 7 minutes with mixing every 2 minutes (plate was placed 3-4 cm away from the bulb). The irradiated cells were then transferred to cold tubes and spun at 330 x g for 4 minutes to pellet. Crosslinked RNA was isolated using TRIzol (Life Technologies) followed by two chloroform extractions and isopropanol precipitation, with the final pellet dissolved in 50 µL water. The purified RNA was fragmented in a 50 µL reaction containing 20-50 µg RNA in 1X fragmentation buffer (Thermo Fisher, AM8740) and heated at 70°C for 2 minutes, at which point 2 µL stop solution (Thermo Fisher, AM8740) was added. The fragmented RNA was then purified using acid phenol/chloroform and the final pellet was dissolved in 100 µL water. The purified RNA was dephosphorylated by adding 1ul 10X cut smart buffer, 1ul rSAP (NEB) to 8ul RNA and incubated at 37°C for 30 minutes, then inactivated at 65°C for 10 minutes. The RNA was then 5'phosphorylated by adding 4ul 10X cut smart buffer, 2.5ul 100 mM DTT, 2.5µl 100 mM ATP, 2.5µl RNAsin (Promega), 2.5µl T4 PNK (NEB), and 26µl H2O. The reaction was incubated at 37°C for 30 minutes and inactivated at 65°C for 10 minutes. The RNA was then purified using the Zymogen RNA clean-up kit. 50 µL of purified RNA was then mixed with 5µl 10X buffer T4 RNA ligase buffer, 2.5µl 100 mM ATP, 2.5µl RNAsin (Promega), 3µl T4 RNA ligase I (NEB), and 25µl H2O. Reactions were incubated overnight at 16°C. SuperScript III (Invitrogen) was used for cDNA synthesis, and NQO1_3utr_RT was used for priming (see Table 1), and samples without reverse transcriptase were used as controls in the subsequent qPCR.

***Crosslinking followed by immunoprecipitation and quantitative RT-PCR (CLIP-qPCR)***
Biological replicates of NQO1-AS knockdown and control MDA-LM2 cells were crosslinked with 400mJ/cm$^2$ 254nm UV. Crosslinked cells were lysed on ice with lysis buffer (100mM Tris pH 7.5, 1% SDS, 1mM EDTA) supplemented with SuperaseIn and 1X protease inhibitors. The lysate was then clarified by spinning at 14,000 x g at 4°C for 10 minutes. The clarified lysate was transferred to protein A dynabeads conjugated to anti-HNRNPC (Santa Cruz, sc-32308) and rotated end-over-end at 4°C for 2 hours. The beads were then washed 1X with high-stringency wash buffer (15mM Tris pH 7.5, 5mM EDTA, 1% Triton X-100, 1% sodium deoxycholate, 0.001% SDS, 120mM NaCl, 25mM KCl), 1X with high-salt wash buffer (15mM Tris pH 7.5, 5mM EDTA, 1% Triton X-100, 1% sodium deoxycholate, 0.001% SDS, 1M NaCl), and 1X with 1X PBS. The immunoprecipitated protein-RNA complexes were then treated with proteinase K (Ambion) in PK reaction buffer (100mM Tris pH 7.5, 100mM NaCl, 1mM EDTA, 0.2% SDS) for 45 minutes at 55°C with intermittent mixing (900rpm 15 seconds/45 seconds rest). The RNA was then extracted with acid phenol-chloroform and ethanol precipitated overnight at -20°C. The purified RNA was then used for quantitative RT-PCR as described above.

***Chromatin immunoprecipitation followed by quantitative RT-PCR (ChIP-qPCR)***
Biological replicates of MDA-Parental and MDA-LM2 cells were crosslinked with 1% paraformaldehyde for 10 minutes at room temperature. The formaldehyde was quenched with 1M glycine (200mM final concentration). The cells were then washed with PBS, scraped from the plate, and collected by centrifugation. Next, the cells were lysed with ChIP lysis buffer (50mM HEPES-KOH pH 7.5, 140mM NaCl, 1mM EDTA pH 8, 1% Triton X-100, 0.1% sodium deoxycholate, 0.1% SDS, 1X protease inhibitors) and the DNA was fragmented by sonication (10 cycles on high, 30 seconds on/30 seconds off, Diagenode Bioruptor UCD-200). An aliquot of the fragmented DNA was removed for size analysis, and the remainder of the sample was diluted 10-fold in RIPA buffer (50mM Tris-HCl pH 8, 150mM NaCl, 2mM EDTA pH 8, 1% NP-40, 0.5% sodium deoxycholate, 0.1% SDS, 1X protease inhibitors) and transferred to protein A dynabeads conjugated to anti-CTCF (Abcam) and rotated end-over-end at 4°C for 3 hours. The beads were then washed once in low salt wash buffer (20mM Tris-HCl pH 8, 150mM NaCl, 0.1% SDS, 1% triton X-100, 2mM EDTA,), once in high salt wash buffer (20mM Tris-HCl pH 8, 500mM NaCl 0.1% SDS, 1% Triton X-100, 2mM EDTA), and once in LiCl wash buffer (10mM Tris-HCl pH 8, 0.25M LiCl, 1% NP-40, 1% sodium deoxycholate, 1mM EDTA). The immunoprecipitated protein-DNA complexes were eluted in elution buffer (1% SDS, 100mM $NaHCO_3$), treated with 2µl RNase A (Thermo Fisher) overnight at 65°C, and treated with 2µl proteinase K (Thermo Fisher) for 1 hour at 60°C. The DNA was purified using a PCR purification kit (Zymogen) and used as a template for qPCR as described above.

**In vitro *proliferation***

In vitro cancer cell proliferation assays were performed by seeding $5 \times 10^4$ cells at day 0 and then counting them in triplicate on day 3 and 5. The slope of the best-fit line between the log of cell counts and days is the reported proliferation rate ($\log Nt = \log N0 + rt$ where t is the time in days and r the proliferation rate per day).

***Animal Studies***

All animal studies were performed according to University of California San Francisco IACUC guidelines. In all cases, seven- to twelve-week-old age-matched female NOD/SCID gamma mice (Jackson Labs) were used.

***Metastatic lung colonization***

Metastatic lung colonization assays were performed by injecting cancer cells stably expressing luciferase into mice via tail vein ($5x10^4$ – $2.5x10^5$ cells per mouse for MDA-Par and MDA-LM2? and $1x10^5$ cells per mouse for HCC1806 cells). In vivo bioluminescence was measured by retro-orbital injection of luciferin (Perkin Elmer) followed by imaging with an IVIS instrument (Perkin Elmer). At the endpoint, the lungs were extracted, fixed with PFA, and subjected to hematoxylin and eosin (H&E) staining.

**In vivo *primary tumor growth***

Orthotopic tumor growth assays were performed by injecting $2.5 \times 10^5$ cells resuspended in 50 µl PBS mixed with 50 µL Matrigel into the mammary glands of female NSG mice using a 28-gauge needle. Tumor volume was assessed using calipers to measure the tumor length (L) and width (W) every 2 days, and calculated using the formula πLW2/6. The experimental end point was reached once tumors reached a volume of 500 $mm^3$.

***ROS measurements***

NQO1 knockdown and control MDA-LM2 and HCC1806-LM2 cells were seeded at a density of $5x10^5$/well in 6-well plates. The following day, reactive oxygen species were measured using the CellROX Green kit (Invitrogen) per the manufacturer's instructions. The cells were then returned to normal growth media and allowed to recover in the incubator for 1 hour. They were then treated with 2 mM tert-Butyl hydroperoxide (TBHP) for 30 minutes and assayed again using the CellROX Green kit (Invitrogen).

***TBHP and $H_2O_2$ sensitivity assays***

NQO1 knockdown and control MDA-LM2 cells were seeded at a density of $2x10^5$/well in 6-well plates. The following day cells were treated with either 0.5mM, 1mM, or 1.5mM $H_2O_2$ or 50uM, 100uM or 150uM TBHP. Cells were counted after 24 hours of treatment. This experiment was repeated in the HCC1806-LM2 background and with MDA-BoM-1833, MDA-BrM2-831, MDA-LM2 and MDA-231-TGL cells.

**In vitro *ferrostatin-1 assay***

$2x10^5$ NQO1 knockdown, NQO1-AS knockdown, and control MDA-LM2 cells were seeded in 6 well plates in media containing 1 µM ferrostatin-1 or DMSO as vehicle control. After 24 hours, the cells were treated with 100uM TBHP or water vehicle. Cells were counted after 24 hours.

**In vitro *liproxstatin-1 assay***

$5x10^3$ NQO1 knockdown and control MDA-LM2 cells were seeded in 96 well plates. The next day cells were treated with 1 µM liproxstatin-1 or DMSO vehicle for 1 hour followed by 100uM liproxstation-1 treatment. After 24 hours, the number of viable cells was determined using the CellTiter-Glo Luminescent Cell Viability Assay (Promega) according to the manufacturer's protocol.

***Metabolomics***

Specialized media was used for cell culture in metabolomics experiments to enable mass spectroscopic analysis of cellular metabolites. For each experiment, half of the cells were grown in "$H_2O$ media", which contained 1X DMEM powder (ThermoFisher #L80677054), 10% dialyzed FBS, 100 units/mL penicillin, 100 µg/mL streptomycin, 1 µg/mL amphotericin, 2 mM L-glutamine, 25 mM glucose, and 40 mM $NaHCO_3$ dissolved in dialyzed $H_2O$. The other half of the cells were grown in "$D_2O$ media", which contained the same components dissolved in 50% dialyzed $H_2O$ and 50% $D_2O$. NQO1 knockdown and control MDA-LM2 or HCC1806-LM2 cells were seeded at a density of $2x10^5$/well in 6-well plates. The following day, cells were treated with 50 µM TBHP or left untreated. 24 hours post-treatment lysates were harvested by adding 400 µL chilled extraction buffer (40:40:20 acetonitrile:methanol:water + 0.5% v/v formic acid) to each well, incubating for 20-40 seconds at room temperature, and quenching with 44 µL neutralization buffer (15% $NH_4HCO_3$ in water). Lysates were transferred to pre-chilled 1.5 mL tubes and frozen at -80 C. Metabolomic profiling was performed by the Rabinowitz lab as previously published[2].

**In vitro *combined drug treatment***

$1.5x10^5$ MDA-Par cells were seeded in 6-well plates and treated the following day with 3uM erastin or DMSO vehicle. 20 hours later, the cells were treated with 15uM rucaparib or DMSO vehicle. 2 hours later, the cells were treated with 1, 2, 3, 4 uM β-lapachone or DMSO vehicle. RNA was then extracted with Quick-RNA Microprep Kit (Zymo Research), and NQO1 mRNA levels were assayed by qPCR.

**In vivo *combined drug treatment***

For in vivo drug treatment studies, $5x10^5$ MDA-Par cells were injected via tail vein. Mice were then injected with Rucaparib (15mg/kg, ip, Sigma Aldrich) or saline control and β-lapachone (22 mg/kg, retroorbital, Sigma Aldrich) or HPβCD vehicle (600 mg/kg, Sigma Aldrich). Erastin (10 mg/kg, ip, Sigma Aldrich) or saline control was injected in the appropriate cohorts. β-lapachone and erastin injections were repeated daily for five days. Lungs were extracted at endpoint and stained with H&E.

***Immunohistochemistry***

Tissue microarrays were obtained from University of Virginia CHTN. After deparaffinization by incubation in two baths of xylene for ten minutes each, the slides were then rehydrated by sequential incubation with 100%, 95%, 80% and 60% ethanol for 5 minutes each. The slides were then rinsed with distilled water 3X for 3 minutes each. Antigen retrieval was done by placing the slides in boiling Tris-EDTA buffer, pH 9.0, and allowed to sit for 35 minutes. The slides were then rinsed 3X with 1X PBS for 3 minutes each, and placed in 3% $H_2O_2$ for 10 minutes to quench endogenous peroxidase activity. The slides were rinsed 3X with 1X PBS for 3 minutes each, and then blocked with 400 ul of 5% milk, diluted in 1X PBST, at room temperature for 1 hour. The slides were then incubated with 400 ul anti-NQO1 antibody (Proteintech 11451-1-AP), at a dilution of 1:200 in 1X PBS overnight at 4°C. The next morning, the slides were rinsed 3X with 1X PBST for 3 minutes each, and then incubated with 400 ul biotinylated secondary antibody (Vector Labs BA-1000), diluted in 1X PBST, at room temperature for 30 minutes. The slides were then washed 3X with 1X PBST for 5 minutes each. Staining was done with Vectastain ABC HRP Kit (Peroxidase, Goat IgG, Vector Labs PK-4005). The ABC reagent was prepared and left at room temperature for 30 minutes, according to manufacturer's instructions. The slides were incubated with 400 ul of the prepared ABC reagent at room temperature for 30 minutes, and then washed 3X with 1X PBST 5 minutes each. The slides were then incubated with ImmPACT DAB Peroxidase (HRP) Substrate (Vector Labs SK-4105) until developed, and afterwards washed in $dH_2O$ 2X for 5 minutes each. The slides were dehydrated by sequential incubation in 60%, 80%, 95%, and 100% ethanol for 5 minutes each, and then incubated in two baths of xylene for 2 minutes each. The slides were air dried and scanned.

#### *Apoptosis assay*

NQO1 knockdown and control MDA-LM2 cells were seeded $4x10^3$ per well in a 96-well plate. The next day, cells were treated with 100 µM TBHP for 24 hours and assayed with the Caspase-Glo 3/7 Assay System (Promega) per the manufactuer's protocol.

#### *IRIS (Identification of antisense RNA species)*

The basis of IRIS is schematized in Fig S1. First, for each gene, we identified the isoform with the longest coding sequence as the representative of that gene. We used the resulting fasta file of sense RNAs as a reference to map GRO-seq reads from MDA-Parental cells (bowtie 2.3.5). Reads mapping to the antisense strand were also tabulated and counted in 500nt increments with a 250nt step. The enrichment of antisense reads in every 500nt window was assessed using logistic regression (Fig S1a). For this the ratio of reads mapping to the 500nt window of interest to the rest of the transcript is compared between the two strands (logASR: log fold-change in anti-sense to sense ratio). Significantly higher presence of reads on the reverse strand is taken as evidence of antisense transcription (logASR>0.5 and FDR<0.01; Fig S1b). Significant neighboring windows were then merged using (Bedtools v2.28.0). The beginning of the first read and end of the last read were used to refine the two ends of the identified antisense RNA species, and both logASR and FDR were re-calculated and the same thresholds were applied. The transcriptomic coordinates of the resulting antisense annotations were converted to genomic coordinates. This process was repeated for a 'background' set of loci which were selected for the absence of any antisense transcript enrichment (logASR~0 and FDR>0.5).

As an independent measure of antisense transcript activity, we asked whether there was evidence of POLR2A binding in or upstream of the antisense of RNA species (above background) based on ENCODE ChIP-seq data. For this, we downloaded narrowPeak bed files (pre-IDR) for all available samples (67 samples total). We used the background loci from above to generate a null distribution for number samples showing POLR2A signal at each locus. We used this distribution to perform outlier analysis on our annotated antisense RNA species. We selected those species that were at least one inter-quantile range above the background median (i.e. median + IQR; Fig S1c). 308 antisense loci passed this filter.

Finally, we used stranded RNA-seq to further validate anti-sense transcription at these loci. This analysis was performed similar to GRO-seq by calculating logASR and the associated *p*-value and FDR. 262 loci with logASR>1 and FDR<1e-5 were defined as our final antisense RNA annotation. Of these 262, 58 were previously annotated antisense RNAs.

#### *Other computational tools*

Salmon v0.14.1 was used to quantify RNA-seq data for both custom sequences or annotated human transcriptomes (gencode v28).

### Figures

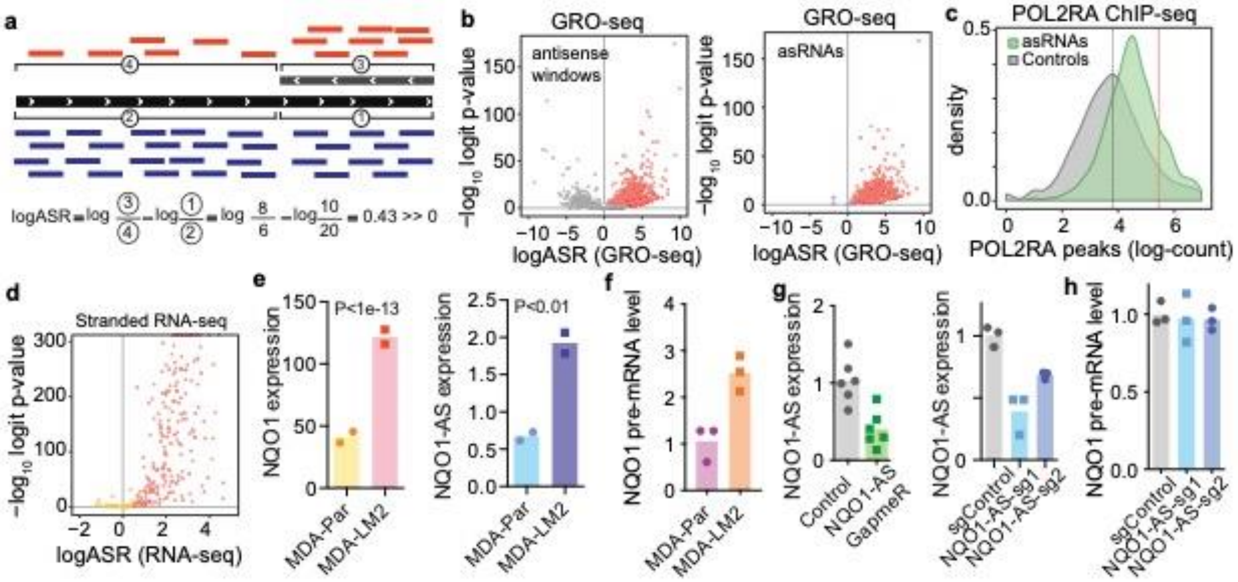


**Figure S1**. **Discovery and annotation of antisense RNAs.** (a) Schematic showing sense and antisense read distribution in IRIS and the calculation of logASR. (b) Volcano plots showing windows of high antisense activity (left) and annotated asRNAs (right) based on GRO-seq data. (c) Prevalence of asRNAs and negative controls in POL2RA ChIP-seq data from ENCODE. Red line represents threshold above which there is strong evidence for RNA Pol II binding across multiple cell lines. (d) Volcano plot showing logASR distribution from stranded RNA-seq in MDA-Par and MDA-LM2 cells. (e) Relative NQO1 (left) and NQO1-AS (right) expression in MDA-Par and MDA-LM2 cells measured by RNA seq. N = 2. (f) Relative NQO1 pre-mRNA level in MDA-Par and MDA-LM2 cells measured by qPCR. N = 3. (g) (Left) Relative NQO1-AS level in control and NQO1-AS GapmeR treated MDA-LM2 cells as measured by qPCR. N = 6. (Right) Relative NQO1-AS level in control cells and cells with CRISPRi-mediated knockdown of NQO1-AS measured by qPCR. N = 3. (h) Relative NQO1 pre-mRNA level in control cells and cells with CRISPRi-mediated NQO1-AS knockdown measured by qPCR. N = 3.

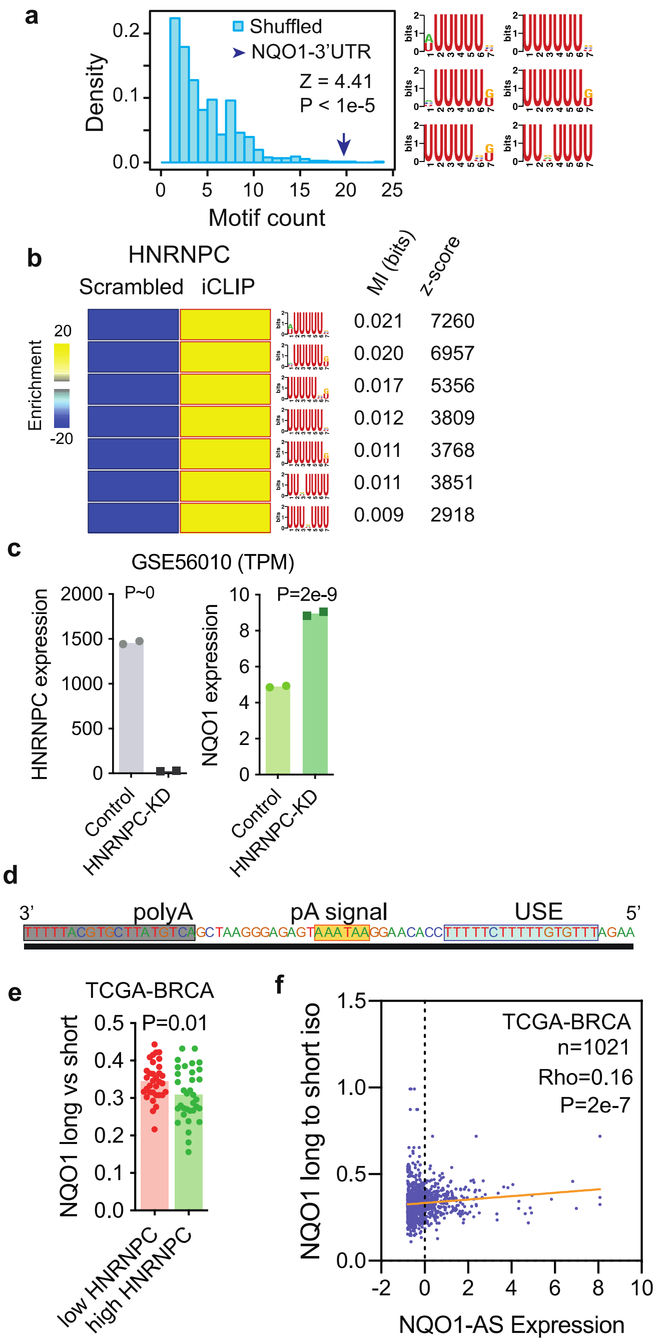


**Figure S2. NQO1-AS binding masks HNRNPC binding sites which modulates polyA site selection.** (a) U-rich motif density in NQO1 3'UTR region complementary to NQO1-AS (purple arrow) relative to scrambled sequences with the same dinucleotide frequency (blue). (b) Heat map showing U-rich motif enrichment in HNRNPC iCLIP data relative to scrambled sequences. (c) Relative HNRNPC (left) and NQO1 (right) expression in HNRNPC knockdown and control (GSE56010; HEK293T) cells measured by RNA-seq. N = 2. (d) Region of NQO1 3'UTR highlighting two canonical polyadenylation sites. (e) Relative long vs short NQO1 isoform ratio in cells with high or low HNRNPC expression from TCGA-BRCA dataset (n=33 and 24 respectively; ~top and bottom 5% of HNRNPC expression values). (f) Correlation between NQO1 long to short isoform ratio and NQO1-AS expression in TCGA-BRCA dataset. N = 1021.

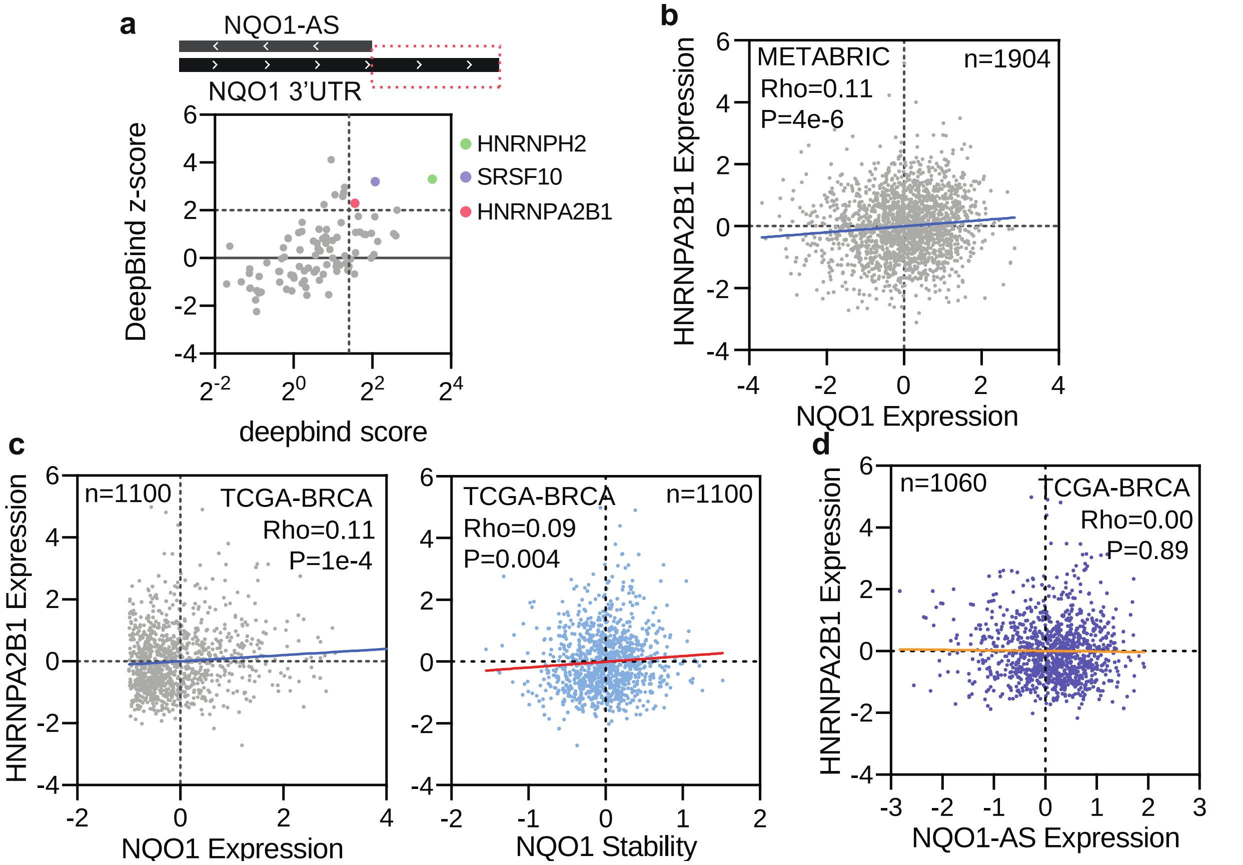


**Figure S3. HNRNPA2B1 binds and stabilizes the long NQO1 isoform and increases NQO1 expression.** (a) DeepBind sequence analysis of NQO1 3'UTR distal region complementary to NQO1-AS. Consensus motifs for HNRNPH2, SRSF10, and HNRNPA2B1 are highlighted. (b) Correlation between NQO1 expression and HNRNPA2B1 expression in METABRIC dataset. N = 1904. (c) Correlation between NQO1 expression (left), NQO1 stability (right) and HNRNPA2B1 expression in TCGA-BRCA dataset. N = 1100. (d) Correlation between NQO1-AS expression and HNRNPA2B1 expression in TCGA-BRCA dataset. N = 1060.

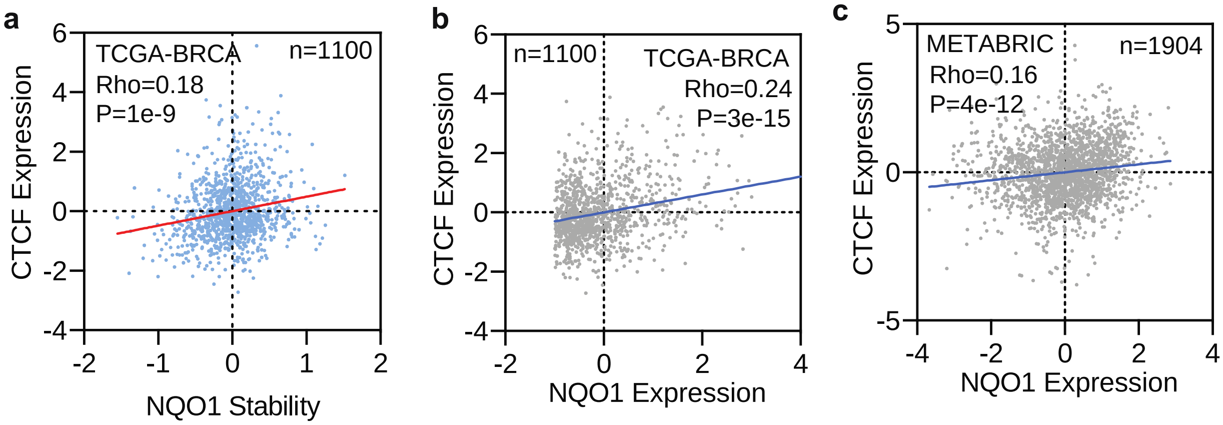


**Figure S4. CTCF binding promotes NQO1-AS transcription in highly metastatic cells.** (a) Correlation between NQO1 stability and CTCF expression in TCGA-BRCA dataset. N = 1100. (b) Correlation between NQO1 expression and CTCF expression in TCGA-BRCA dataset. N = 1100. (c) Correlation between NQO1 expression and CTCF expression in METABRIC dataset. N = 1904.

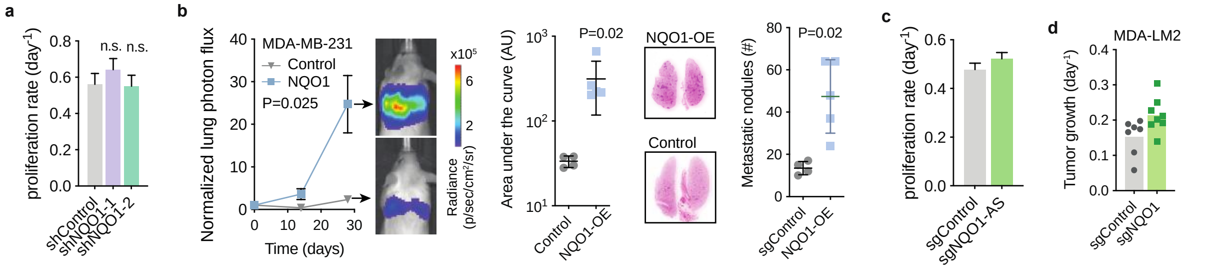


**Figure S5. NQO1 and NQO1-AS promote metastatic lung colonization.** (a) *in vitro* proliferation rate of control and shRNA-mediated NQO1 knockdown MDA-LM2 cells. N = 6. (b) *in vivo* lung colonization assay with MDA-Par control and NQO1 overexpression cells. Bioluminescence over time is shown on the left alongside representative images from each cohort at the end point. Total bioluminescence over the course of the experiment and nodule count at the end point are shown on the right, alongside representative H&E-stained lungs. N = 4. (c) *in vitro* proliferation rate of control and CRISPRi-mediated NQO1-AS knockdown MDA-LM2 cells. N = 6. (d) *in vivo* primary tumor growth assay with control and NQO1 knockdown MDA-LM2 cells. N = 7.

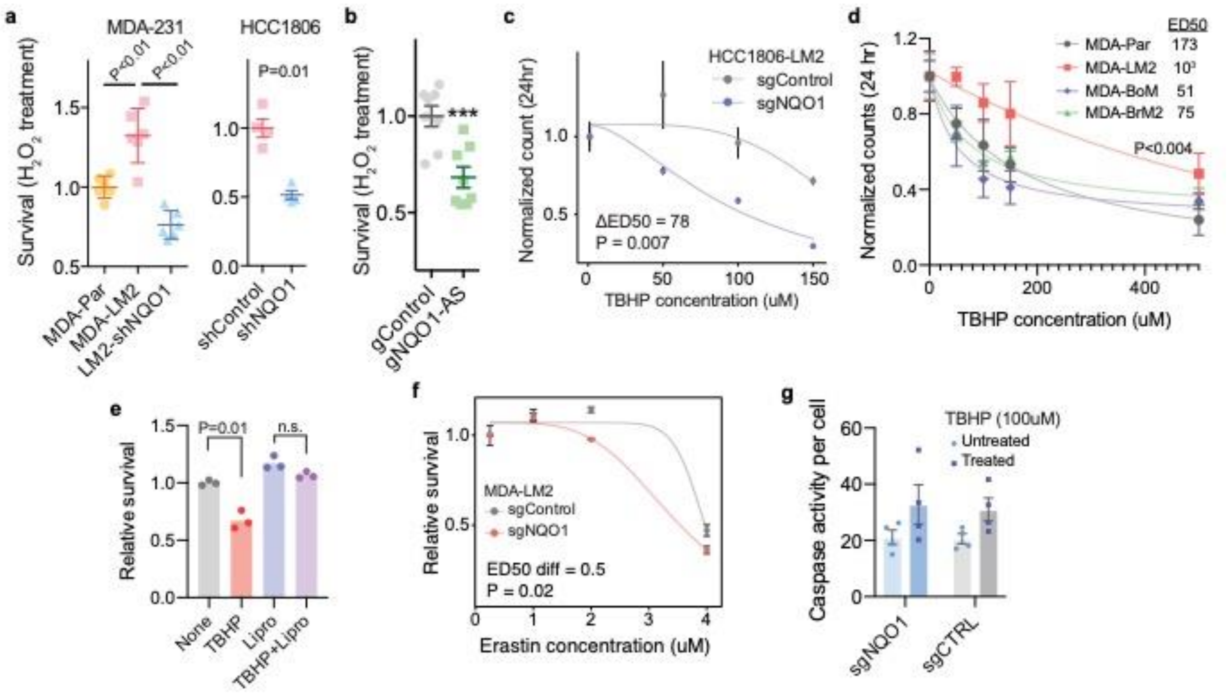


**Figure S6. NQO1 protects cancer cells from ferroptosis.** (a) (Left) Relative survival of MDA-Par, MDA-LM2, and shRNA-mediated NQO1 knockdown MDA-LM2 cells after treatment with $H_2O_2$. N = 6. (Right) Relative survival of HCC1806-LM2 control and shRNA-mediated NQO1 knockdown cells after treatment with $H_2O_2$. N = 4. (b) Relative survival of control and CRISPRi-mediated NQO1-AS knockdown MDA-LM2 cells after treatment with $H_2O_2$. N = 8. (c) Relative sensitivity of HCC1806-LM2 control and NQO1 knockdown cells to TBHP. N = 6. (d) Relative sensitivities of MDA-Par, MDA-LM2, MDA-BoM, and MDA-BrM2 cells to TBHP. N = 3. (e) Relative survival of MDA-LM2 cells treated with TBHP, Liproxstatin-1, TBHP + Liproxstatin-1, or vehicle control. N = 3. (f) Relative sensitivity of MDA-LM2 control and NQO1 knockdown cells to Erastin. N = 3. (g) Relative caspase activity in TBHP treated and untreated NQO1 knockdown and control MDA-LM2 cells. Caspase activity was measured as a proxy for apoptotic cell death using the Caspase Glo 3/7 Assay System from Promega. N = 4.

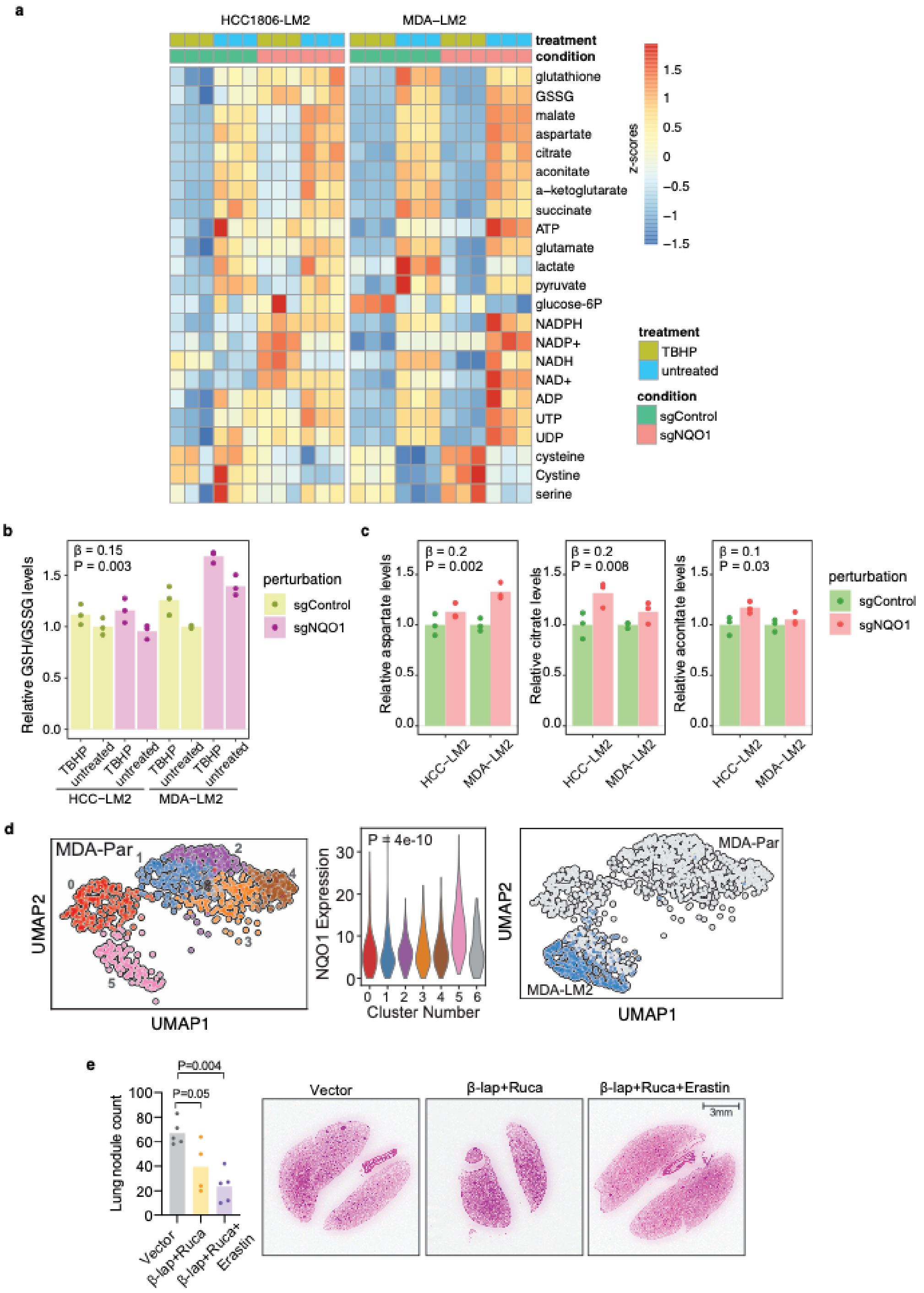


**Figure S7. NQO1 mediates metabolic remodeling in cancer cells.** (a) Heatmap showing metabolic changes in MDA-LM2 and HCC1806-LM2 NQO1 knockdown and control cells after treatment with TBHP. Metabolite levels were measured using LC/MS profiling. (b) Relative NADPH, aspartate, citrate, and aconitate levels in NQO1 knockdown and control MDA-LM2 and HCC1806-LM2 cells. N = 3. (c) UMAP visualization of single cell RNA-seq data from MDA-Par cells (left), NQO1 expression level by cluster (middle), and overlay with MDA-LM2 data (right). (d) End point histologic staining after *in vivo* lung colonization assay with MDA-LM2 cells and subsequent treatment with either β-Lapachone + Rucaparib, β-Lapachone + Rucaparib + Erastin, or vector control. Lung nodule counts for each cohort are shown on the left, and representative images are shown on the right. N = 5.

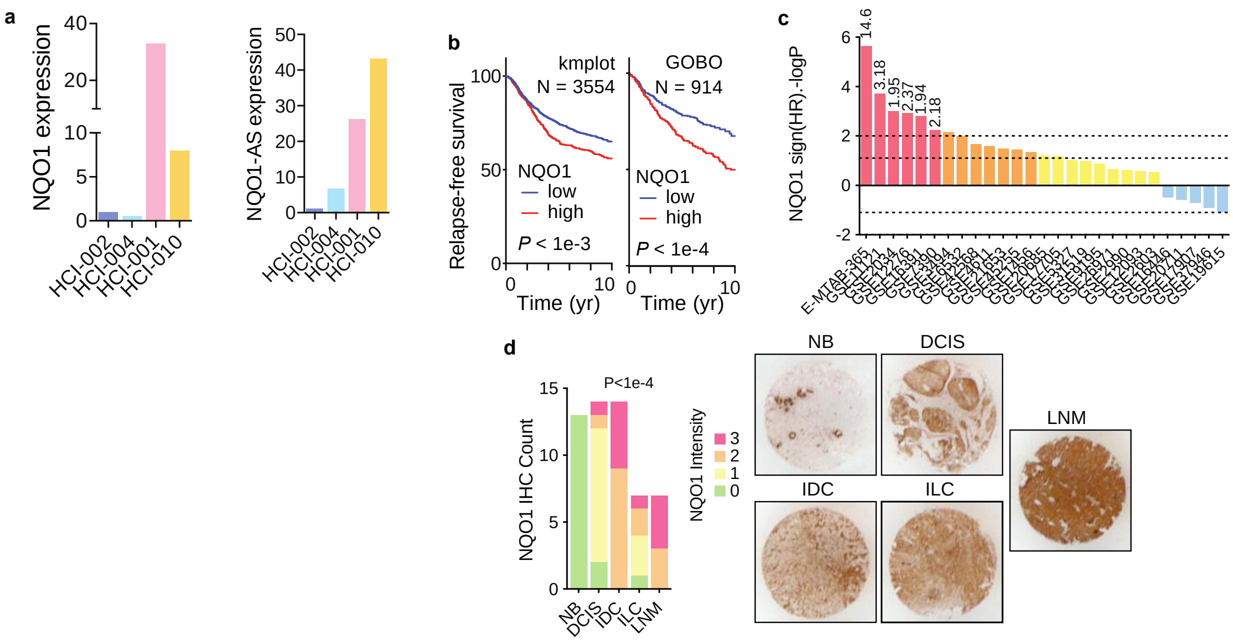


**Figure S8. NQO1-AS and NQO1 expression are associated with metastasis in clinical samples**. (a) Relative NQO1 (left) and NQO1-AS (right) expression in 4 patient-derived xenograft lines as measured by RNA-seq. (b) Kaplan-Meier curve showing relative relapse free survival of patients with tumors expressing high and low levels of NQO1 from kmplot (left, N = 3554) and GOBO (right, N = 914) databases. (c) Distribution of 10-year relapse-free survival *P* values (two-sided log rank test results reported as –logP for positive association and logP for negative) for the correlation of NQO1 expression and clinical outcome in the listed breast cancer datasets. Red bars show associations that pass the statistical threshold, orange bars are trending positive, and blue bars are trending negative. For statistically significant datasets, the hazard ratio is included at the top of the bar. (d) Immunohistochemical staining of NQO1 in a tissue microarray containing non-neoplastic breast tissue (NB), ductal carcinoma in situ (DCIS), invasive ductal carcinoma (IDC), invasive lobular carcinoma (ILC), and lymph node metastases (LNM). Blinded grading of the stain intensity is represented by the bar graph on the left. Representative images of stained specimens are shown on the right.
